# Engineered NKG2C^+^ NK-like T cells exhibit superior antitumor efficacy while mitigating cytokine release syndrome

**DOI:** 10.1101/2024.07.16.603785

**Authors:** Kyle B. Lupo, M. Kazim Panjwani, Sanam Shahid, Rosa Sottile, Clara Lawry, Gabryelle Kolk, Theodota Kontopolous, Anthony F. Daniyan, Smita S. Chandran, Christopher A. Klebanoff, Katharine C. Hsu

**Affiliations:** Human Oncology and Pathogenesis Program, Memorial Sloan Kettering Cancer Center, New York, NY, USA. 10065; Humanitas Clinical and Research Center, 20090, Pieve Emanuele, Italy; Department of Medicine, Memorial Sloan Kettering Cancer Center, New York, NY, USA; Center for Cell Engineering and Department of Medicine, MSKCC, New York, NY, USA. 10065; Department of Medicine, Weill Cornell Medical College, New York, NY, USA. 10065; Parker Institute for Cancer Immunotherapy, MSKCC, New York, NY, USA. 10065; Immunology Program, Sloan Kettering Institute, Memorial Sloan Kettering Cancer Center, New York, NY, USA

## Abstract

Engineered T and NK cell therapies have widely been used to treat hematologic malignancies and solid tumors, with promising clinical results. Current chimeric antigen receptor (CAR) T cell therapeutics have, however, been associated with treatment-related adverse events such as cytokine release syndrome (CRS) and are prone to immunologic exhaustion. CAR-NK therapeutics, while not associated with CRS, have limited in vivo persistence. We now demonstrate that an NK-like TCRαβ^+^ CD8 T cell subset, identified and expanded ex vivo through its expression of the activating receptor NKG2C (NKG2C^+^ NK-like T cells), can be transduced to express a second-generation CD19 CAR (1928z), resulting in superior tumor clearance, longer persistence and decreased exhaustion compared to conventional 1928z CAR^+^ CD8 T cells and 1928z CAR+ NK cells. Moreover, CAR-modified NKG2C^+^ NK-like T cells resulted in significantly reduced CRS compared to conventional CAR^+^ CD8 T cells. Similarly, NKG2C^+^ NK-like T cells engineered with a TCR targeting the NY-ESO-1 antigen exhibit robust tumor control and minimal exhaustion compared to TCR-engineered conventional CD8 T cells. These data establish NKG2C^+^ NK-like T cells as a robust platform for cell engineering, and offer a safer, more durable alternative to conventional CAR-T and CAR-NK therapies.

## Introduction

Despite promising advances in chimeric antigen receptor (CAR)- and T cell receptor (TCR)-engineered T cell therapeutics for treating hematologic malignancies, major hurdles, including cytokine release syndrome (CRS), neurotoxicity, and antigen-induced exhaustion, limit safe and efficacious use of engineered T cell therapeutics in many hematologic malignancies and solid tumors ^1, 2, 3, 4, 5^. CRS has been reported in 37% to 93% of patients post-CAR-T treatment across different studies, and immune effector cell-associated neurotoxicity syndrome (ICANS) has been reported in 40% of patients receiving CAR-T therapies ^3, 4^. Patients receiving CAR-T therapies experience a non-relapse mortality rate of approximately 5.0%, with greater than 20% of patients experiencing high-grade CRS and/or ICANS requiring intensive monitoring and medical intervention^3, 4^. Further, continuous antigen signaling and upregulation of inhibitory receptors, such as PD-1, contribute to the exhaustion and eventual failure of CAR-engineered T cells in many settings ^1^. As a consequence, patients with B cell malignancies have a high incidence of relapse post-treatment (14% for all non-Hodgkin’s lymphomas^6^, 23% for diffuse large B cell lymphoma^7^, and 30-45% for chronic lymphocytic leukemia^8^), spurring the need for novel long-lasting exhaustion-resistant therapeutic options.

The emergence of CAR-modified natural killer (NK) cell therapies has offered new hope to patients, given their lack of associated CRS, neurotoxicity, and exhaustion^9^. A recent clinical trial utilizing CD19-targeted umbilical cord-derived CAR-NK cells demonstrated no notable toxicities, including CRS, neurotoxicity, or graft-versus-disease, while reporting 30 and 100 day overall response rates of 48.6% each^10^. Poor in vivo persistence following transfer, however, remains a major limitation to their therapeutic potential^11^.

Though innate activity from T cells has been long-described in subsets, such as γδ T cells, invariant NKT cells, and MAIT cells, we and others have identified and characterized a wider variety of populations of TCRαβ^+^ CD8 T cells that straddle the boundary of innate and adaptive immunity, exhibit broader TCR diversity, and express cell surface markers traditionally expressed on NK cells, including, CD56, KIR, NKp30, and NKG2 molecules^12, 13, 14, 15, 16, 17, 18, 19, 20, 21, 22, 23^. We recently described one specific NK-like CD8 T cell population that is frequently found in human cytomegalovirus (HCMV) seropositive individuals, is readily identifiable by NKG2C surface expression, and is phenotypically and transcriptionally remarkably similar to NK cells, resulting in its expression of a full array of NK receptors, including CD56, inhibitory KIRs, NKp30, NKp46, 2B4, CD16 and NKG2D^17^. Beyond their striking phenotypic and transcriptional resemblance to NK cells, these NK-like CD8 T cells interact with HLA-E through NKG2C and display NK-like recognition and cytotoxicity toward malignant and virally infected cells. Fundamental to this T cell population’s assumption of an NK-like identity is its loss of BCL11B^17^, the transcription factor recognized to be the master regulator of T cell fate^24, 25, 26, 27^. NKG2C^+^ NK-like T cells are further distinguished from other NK-like T cell subsets by their exclusive expansion in HCMV individuals, their oligoclonality and their restriction by HLA-E^5, 12, 14, 15, 17, 18, 19, 21^. The endogenous TCR is dispensable for the anti-tumor activity of these cells, and although NKG2C contributes to tumor targeting via interaction with HLA-E on the target cell surface, the cells still mediate innate-like tumor targeting through other NK activating receptors when NKG2C is blocked^17^. While NKG2C is therefore one of an array of activating NK receptors on the NK-like CD8 T cell population, it is a useful marker of the NK-like T cell population, permitting easy identification, isolation, and selective expansion, features facilitating its use as a cell therapy over other T cell populations. Recognizing that NKG2C^+^ NK-like T cells possess phenotypic and functional qualities of both T and NK cells, we hypothesized that they might therefore offer important advantages over traditional CAR-T and CAR-NK cell therapies, particularly regarding side-effect profile and in vivo persistence, respectively.

We have tested the utility of NKG2C^+^ NK-like T cells as a cell therapy platform, engineered to target both hematologic and solid tumors. We find that these cells can be robustly expanded using an HLA-E^+^ K562-based feeder cell platform and efficiently modified via retroviral transduction. CAR-modified NKG2C^+^ NK-like T cells demonstrate superior in vitro and in vivo cytotoxicity and persistence against B cell malignancies over CAR-modified conventional (NKG2C^-^NKG2A^-^KIR^-^) CD8 T cells and NK cells. Importantly, CAR-modified NKG2C^+^ NK-like T cells exhibit significantly reduced CRS, do not upregulate traditional T cell exhaustion markers in response to tumor targets and demonstrate longer persistence in comparison to conventional CAR^+^ conventional CD8 T cells in murine xenograft models. Further, while NKG2C^+^ NK-like T cells modified with a transgenic NY-ESO-1 TCR exhibit similar anti-tumor effector functions to conventional TCR-engineered CD8 T cells, they demonstrate enhanced in vivo persistence and decreased expression of exhaustion markers. NKG2C^+^ T cells exhibit advantageous features of both NK cells and conventional CD8 T cells, ultimately resulting in its superiority to each as a safer and more efficacious effector cell population for anti-tumor therapy.

## Results

### CAR-modified NKG2C^+^ NK-like T cells demonstrate superior in vitro killing of CD19^+^ targets when compared to conventional CD8 T cells

The NKG2C^+^ NK-like CD8 T cell population naturally expands in ∼40% of HCMV-seropositive individuals. We identified individuals with NKG2C^+^ NK-like T cells and compared the suitability of this population as an engineered cell population to conventional CD8 T cell and NK cell populations. We isolated NKG2C^+^ NK-like T cells, conventional CD8 T cells, and NK cells from each donor and transduced each population with a CD19-targeted CAR construct consisting of an anti-CD19 single chain variable fragment (scFv), CD28 hinge, transmembrane, and intracellular domains, as well as a CD3z intracellular signaling domain (1928z) with an mCherry reporter, all under a single promoter (*Figure 1A-B, S1*). Retroviral transduction of NKG2C^+^ NK-like T cells yielded high expression of 1928z^+^ cells (70.90% ± 5.80%), superior to that of conventional CD8 T cells (44.11% ± 7.23%) and NK cells (42.94% ± 6.21%) (*Figure 1A-B*). Further, NKG2C^+^ NK-like T cells with or without CAR-modification robustly expanded ex vivo when cocultured with an irradiated HLA-E-expressing K562-based feeder platform (*Figure S2*). Following incubation with CD19^+^ NALM6 target cells, 1928z CAR-expressing T and NK cells were immunophenotyped for expression of activating receptors and exhaustion markers. The 1928z^+^ NKG2C^+^ NK-like T cells significantly upregulated expression of the activating receptor 4-1BB compared to 1928z^+^ conventional CD8 T cells and 1928z^+^ NK cells following stimulation with NALM6 target cells. Notably, they did not upregulate PD-1 and TIM-3, however, while 1928z^+^ conventional CD8 T cells expressed high levels of these exhaustion markers (*Figure 1C*). LAG-3 was not upregulated in either group. Unmodified cells co-incubated with NALM6 target cells were also evaluated, where conventional CD8 T cells expressed higher 4-1BB and LAG-3 than NKG2C^+^ NK-like T cells and NK cells, and NKG2C^+^ NK-like T cells expressed lower levels of TIM-3 than conventional CD8 T cells and NK cells at baseline (*Figure S3*). Unlike the CAR-modified cell groups, no change in marker expression was observed after target stimulation.

**Figure 1.**
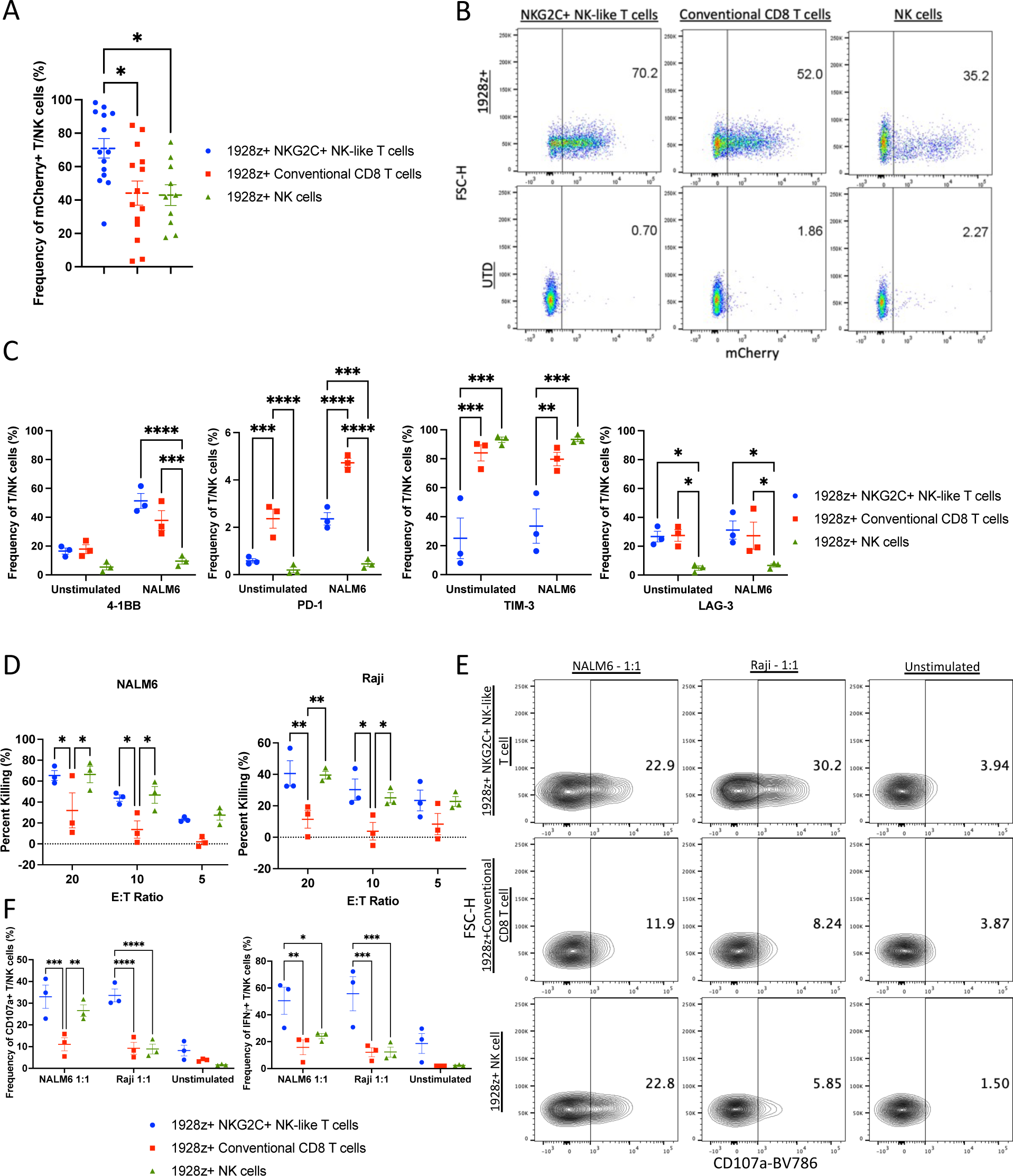
CAR^+^ NKG2C^+^ NK-like T cells mediate robust effector functions against CD19^+^ tumor targets. (A) CAR expression and (B) representative pseudocolor plots showing transduction efficiency measured via mCherry expression via flow cytometry (n=14 independent experiments). (C) Flow cytometry phenotyping of CAR^+^ T and NK cells with and without 8-hour stimulation with CD19^+^ NALM6 target cells (n=3 independent experiments). (D) Cytotoxicity of CAR^+^ T and NK cells against CD19^+^ NALM6 and Raji target cells (n=3 independent experiments). (E) Representative contour plots of CD107a expression of CAR^+^ T and NK cells following 4-hour coculture with CD19^+^ NALM6 and Raji target cells. (F) CD107a and IFN-γ expression of CAR^+^ T and NK cells following 4-hour coculture with CD19^+^ NALM6 and Raji target cells (n=3 independent experiments). Data are presented as mean values +/- SEM. Statistical analysis was performed by one-way or two-way ANOVA.

We characterized 1928z^+^ NKG2C^+^ NK-like T cells for cytolysis, degranulation, and IFN-γ secretion in response to NALM6 leukemia and CD19^+^ Raji lymphoma cell lines in vitro. 1928z^+^ NKG2C^+^ NK-like T cells demonstrated efficient cytolysis of both NALM6 and Raji target cells, superior to that of 1928z^+^ conventional CD8 T cells and similar to that of 1928z^+^ NK cells (*Figure 1D*). The innate-like tumor cytolysis capacity of these NK-like T cells was underscored by the cytolysis of NALM6 targets by non-engineered NKG2C^+^ NK-like T cells and NK cells, but not by autologous conventional CD8 T cells (*Figure S4).* Further, in cytotoxicity assays against CD19-KO NALM6 tumor targets, both 1928z^+^ NKG2C^+^ NK-like T cells and 1928z^+^ NKG2C^+^ NK cells efficiently lysed target cells, highlighting not only the innate-like tumor capacity of 1928z^+^ NKG2C^+^ NK-like T cells but also their resistance to antigen-escape mechanisms (*Figure S5*). Finally, 1928z^+^ NKG2C^+^ NK-like T cells demonstrated enhanced degranulation and IFN-γ production, compared to 1928z^+^ conventional CD8 T cells and 1928z^+^ NK cells in response to the CD19^+^ target cell lines (*Figure 1E-F*). Together, the data show the superior in vitro cytotoxicity of CAR-modified NKG2C^+^ NK-like T cells compared to their conventional CD8 counterparts.

### CAR^+^ NKG2C^+^ NK-like T cells are superior to CAR^+^ conventional CD8 T cells for in vivo tumor killing

Immunodeficient NRG mice engrafted with NALM6 tumors were treated with 1928z^+^ or untransduced NKG2C^+^ NK-like T cells, conventional CD8 T cells, or NK cells and evaluated for tumor burden, and survival (*Figure 2A)*. We previously showed that like NK cells, NKG2C^+^ NK-like T cells exhibit high expression of *IL2RB*, suggesting that NKG2C^+^ NK-like T cells may be highly responsive to IL-15^17^. Indeed, mice receiving 1928z^+^ NKG2C^+^ NK-like T cells and recombinant human IL-15, which is not cross reactive between mice and humans, demonstrated a dramatic reduction in tumor progression compared to all other treatment groups (*Figure 2B-C, S6A-B*). Accordingly, 1928z^+^ NKG2C^+^ NK-like T cells supplemented with IL-15 significantly improved survival of NALM6 tumor-bearing mice over those receiving IL-15 and 1928z^+^ conventional CD8 T cells or 1928z^+^ NK cells (*Figure 2D*). Without IL-15, mice receiving 1928z^+^ NKG2C^+^ NK-like T cells still exhibited improved survival, albeit more modest, over those receiving 1928z^+^ conventional CD8 T cells or 1928z^+^ NK cells (*Figure S6C*). No significant differences in tumor progression or survival were observed between mice treated with the untransduced NKG2C^+^ NK-like T cells, conventional CD8 T cells or NK cells, regardless of IL-15 supplementation (*Figures S7-S8).* Together, these data indicate that 1928z^+^ NKG2C^+^ NK-like T cells not only have superior anti-tumor efficacy to 1928z^+^ conventional CD8 T cells and 1928z^+^ NK cells, but that they are also responsive to IL-15, which further enhances efficacy. All subsequent in vivo efficacy studies therefore employed IL-15 supplementation, an accepted supplementation to adoptive CAR-NK therapies^10, 28^.

**Figure 2.**
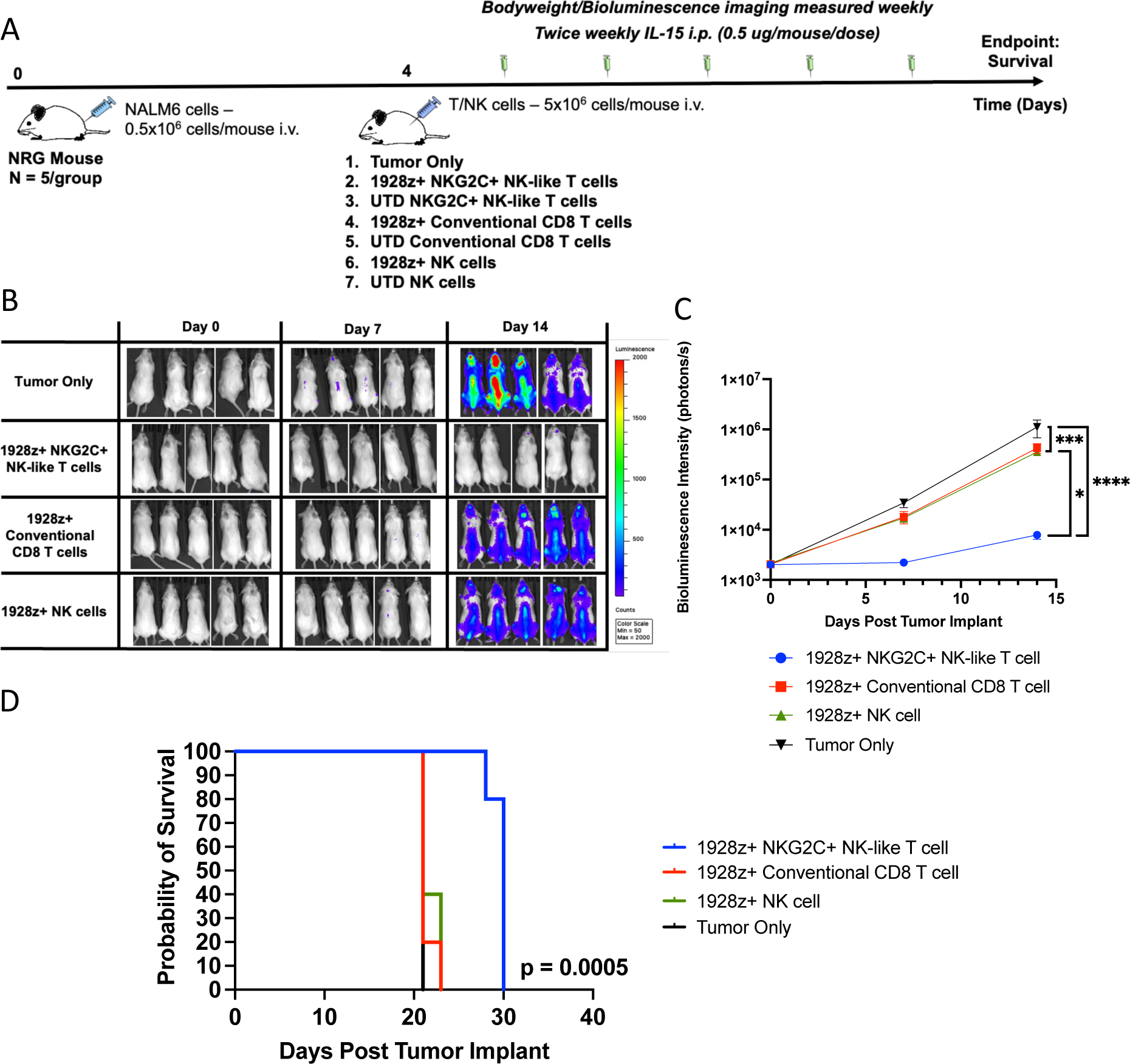
CAR^+^ NKG2C^+^ NK-like T cells mediate enhanced tumor control over CAR^+^ conventional CD8 T cells and CAR^+^ NK cells. (A) Study timeline of NALM6 tumor-bearing mice treated with 1928z CAR-engineered or untransduced (UTD) NKG2C^+^ NK-like T cells, conventional CD8 T cells, or NK cells, with or without IL-15 supplementation. (B) Bioluminescence imaging of mice treated with tumor only, CAR^+^ NKG2C^+^ NK-like T cells, CAR^+^ conventional CD8 T cells, and CAR^+^ NK cells measured throughout the course of the study (n=5 mice/group). (C) Quantification of bioluminescence imaging of mice measured throughout the course of the study (n=5 mice/group). (D) Kaplan-meier survival plot of tumor only, CAR^+^ NKG2C+ NK-like T cell-, CAR^+^ conventional CD8 T cell-, and CAR^+^ NK cell-treated (n=5 mice/group). Data are presented as mean values +/- SEM. Statistical analysis was performed by two-way ANOVA or Kaplan-Meier survival analysis.

### CAR-modified NKG2C^+^ NK-like T cells do not upregulate exhaustion markers associated with T cell dysfunction

Following infusion into tumor-bearing mice, circulating cells were collected at seven days post-treatment, evaluated for persistence and immunophenotyped for exhaustion (*Figure 3, S9-12)*. Mice receiving 1928z^+^ NKG2C^+^ NK-like T cells had higher levels of circulating cells post-treatment than mice receiving either 1928z^+^ conventional CD8 T cells or 1928z^+^ NK cells, with or without IL-15; and mice receiving 1928z^+^ NKG2C^+^ NK-like T cells with IL-15 had the highest persistence of circulating CD8 T cells over all other treatment groups (*Figure 3A, S10A*). Untransduced NKG2C^+^ NK-like T cells also demonstrated higher persistence in mice treated with IL-15 than conventional CD8 T cells and NK cells (*Figure S11)*. The majority of 1928z^+^ NKG2C^+^ NK-like T cells exhibited an effector (CD45RA^+^CD62L^-^CCR7^-^) and effector memory (CD45RA^-^ CD62L^-^CCR7^-^) phenotype, while the majority of 1928z^+^ conventional CD8 T cells exhibited a naïve (CD45RA^+^CD62L^+^CCR7^+^) and central memory (CD45RA^-^CD62L^+^CCR7^+^) phenotype after in vivo activation, with no notable differences observed between groups treated with or without IL-15 (*Figure 3B-C, S10B-C*). These data are consistent with the enhanced effector differentiation of CAR-modified NKG2C^+^ NK-like T cells *(Figures 1-2).* While TEMRA-like, the observed CD45RA^+^CD62L^-^CCR7^-^ phenotype is also consistent with staining of a resting NK cell phenotype, further endorsement of the cell’s NK-like identity^17^. When evaluated for activation markers, both 1928z^+^ NKG2C^+^ NK-like T cells and 1928z^+^ conventional CD8 T cells exhibited higher levels of CD69 and 4-1BB than 1928z^+^ NK cells (*Figure 3D*). Strikingly, despite a high level of engraftment, 1928z^+^ NKG2C^+^ NK-like T cells did not upregulate exhaustion markers PD-1, TIM-3, or LAG-3, in comparison to 1928z^+^ conventional CD8 T cells (*Figure 3D, S10D*). When compared to 1928z^+^ NK cells, 1928z^+^ NKG2C^+^ NK-like T cells exhibited similarly low levels of PD-1, dramatically lower levels of TIM-3 and intermediate levels of LAG-3. Taken together, these data demonstrate that CAR-modified NKG2C^+^ NK-like T cells exhibit enhanced persistence without exhaustion compared with CAR-modified conventional CD8 T cells and NK cell therapies following adoptive transfer into tumor-bearing hosts.

**Figure 3.**
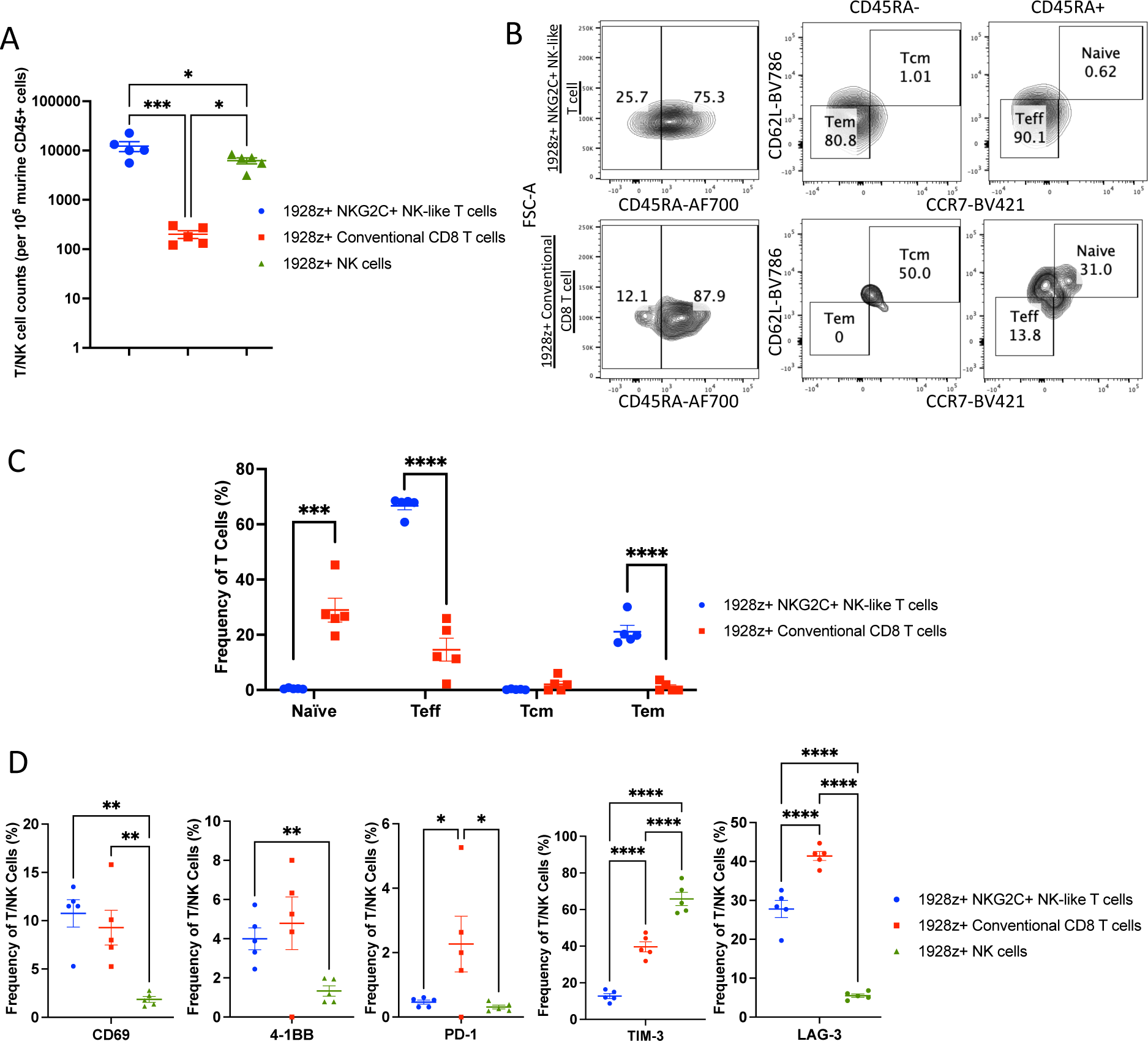
CAR^+^ NKG2C^+^ NK-like T cells demonstrate increased persistence, enhanced activation, and reduced exhaustion over CAR^+^ conventional CD8 T cells and CAR^+^ NK cells. (A) Aggregate quantification, via flow cytometry, of circulating CAR^+^ NKG2C^+^ NK-like T cells, CAR^+^ conventional CD8 T cells, and CAR^+^ NK cells in mice at day 7 post-infusion (n=5 mice/group). (B) Representative contour plots of circulating naïve, Teff, Tem, and Tcm populations among CAR^+^ T cells at day 7 post-infusion measured via CD45RA/CCR7/CD62L flow cytometry staining (C) Aggregate quantification of circulating naïve, Teff, Tem, and Tcm populations among CAR^+^ T cells at day 7 post-treatment measured via CD45RA/CCR7/CD62L flow cytometry staining (n=5 mice/group) (D) Activation and exhaustion marker phenotype of circulating CAR^+^ T and NK cells at day 7 post-infusion (n=5 mice/group). Data are presented as mean values +/- SEM. Statistical analysis was performed by one-way ANOVA or two-tailed unpaired t-test.

### CAR-modified NKG2C^+^ NK-like T cells exhibit reduced CRS compared to conventional CD8 T cells

Traditional T cell therapies have been associated with CRS while NK therapies have not, leading to speculation as to how the NK-like T cell population would behave. We therefore used an established mouse CRS model to test the capacity for CAR^+^ NKG2C^+^ NK-like T cells to induce CRS^3, 4^. In accordance with other studies using this model, CRS was identified through weight loss compared to a tumor only control, an increase in a panel of human and murine cytokines following immune cell administration, and an increase in mSAA-3, the murine equivalent of C-reactive protein, an inflammatory response protein and an early indicator of CRS increased in patients^29, 30^. SCID beige mice, which are deficient for T, B, and NK cells but have a functional myeloid compartment, were implanted with CD19^+^ Raji tumors, and tumors were allowed to engraft for 21 days prior to T cell administration (*Figure 4A*). Mice were then injected with either 1928z^+^ NKG2C^+^ NK-like T cells or 1928z^+^ conventional CD8 T cells and monitored for 72 hours. Mice receiving 1928z^+^ conventional CD8 T cells exhibited a significant loss in bodyweight compared to mice receiving 1928z^+^ NKG2C^+^ NK-like T cells and tumor only controls (*Figure 4B*). Sera collected at 24 hours from mice receiving 1928z^+^ conventional CD8 T cells had significantly higher levels of mSAA-3, compared to the same mice pre-treatment. In contrast, mice receiving 1928z^+^ NKG2C^+^ NK-like T cells did not experience an elevation in mSAA-3 (*Figure 4C*). Sera from mice were assessed for pro-inflammatory cytokines known to correlate strongly with CRS severity, including CXCL1, IL-6, CCL2, G-CSF, IL-3, IFN-g, GM-CSF, and IL-2^29^. Mice receiving 1928z^+^ conventional CD8 T cells exhibited higher levels of hGM-CSF, hIL-3, and hIFN-g in comparison to pre-treatment controls, while no upregulation was observed in mice that received 1928z^+^ NKG2C^+^ NK-like T cells (*Figure 4D*). Both cohorts of mice exhibited equivalently higher levels of IL-2 in comparison to pre-treatment controls (*Figure 4D*). Sera from mice that received 1928z^+^ conventional CD8 T cells also showed elevation in a number of murine cytokines, including mIL-5, mIL-6, mG-CSF, and mCXCL1. Strikingly, these murine cytokine elevations were not observed in mice treated with 1928z^+^ NKG2C^+^ NK-like T cells (*Figure 4E*). No significant differences in serum cytokine levels for hIL-6, hG-CSF, hTNF, mIFNg, mIL-3, mGM-CSF, or mCXCL9 were observed (*Figure S13)*. Finally, murine immune subsets in the blood, bone marrow, spleen, and peritoneal cavity were quantified at 72 hours following T cell treatment to assess CRS-induced expansion of myeloid subsets following T cell administration. Mice receiving 1928z^+^ conventional CD8 T cells displayed a higher frequency of splenic macrophages, monocytes, and dendritic cells (DCs) than mice receiving 1928z^+^ NKG2C^+^ NK-like T cells (*Figure S14-15).* Taken together, the differences in weight loss and human and murine serum cytokine measurements, and splenic infiltration with myeloid cells indicate lower CRS induction with CAR-modified NKG2C+ NK-like CD8 T cells compared to conventional CD8 T cells.

**Figure 4.**
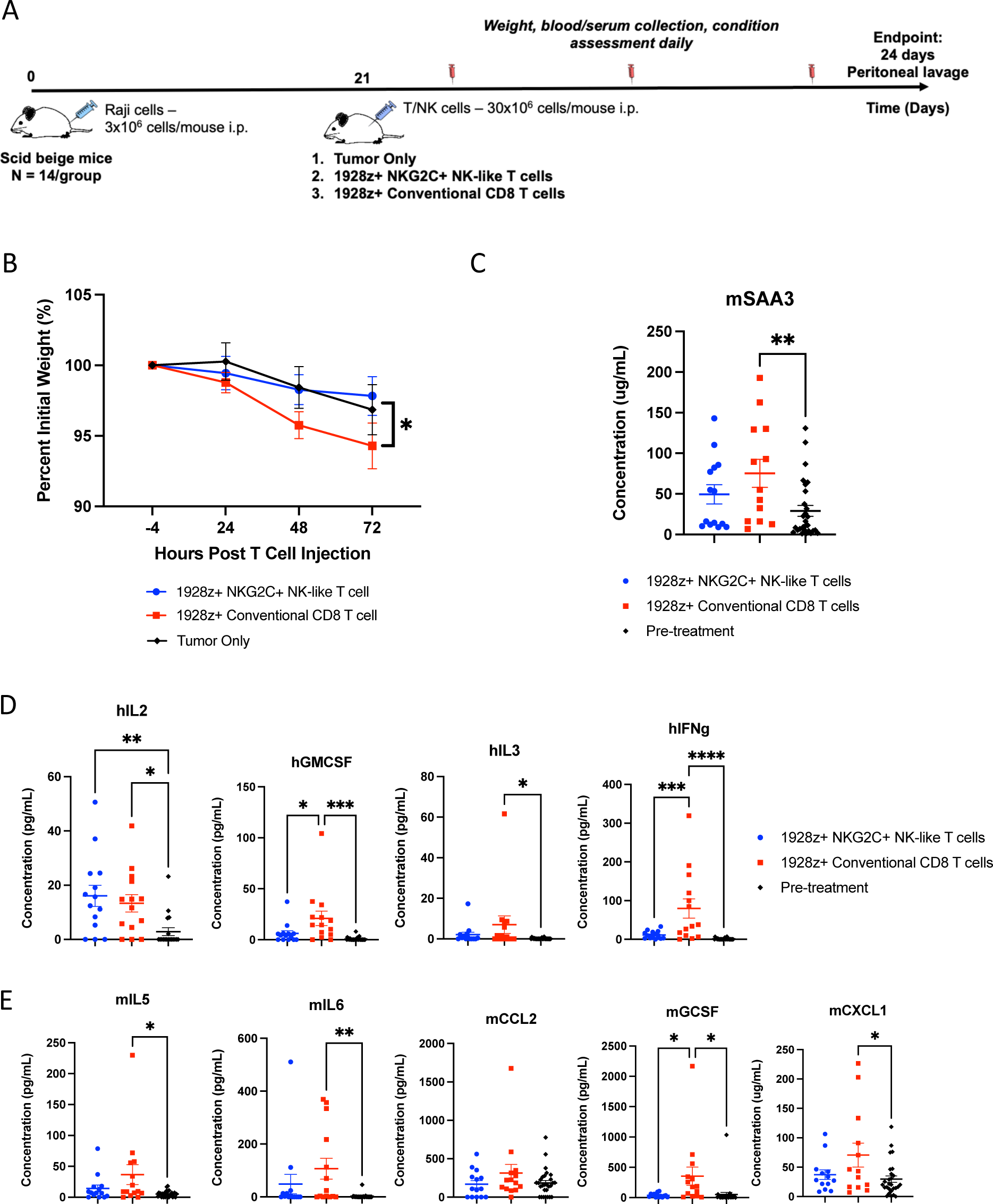
CAR^+^ NKG2C^+^ NK-like T cells do not mediate CRS. (A) Study timeline of Raji tumor-bearing mice treated with or without 1928z CAR-engineered NKG2C^+^ NK-like T cells or conventional CD8 T cells. (B) Bodyweight measurements of mice treated with tumor only, CAR^+^ NKG2C^+^ NK-like T cells, and CAR^+^ conventional CD8 T cells throughout the course of the study (n=14 mice/group). (C) Serum levels of murine SAA-3 from mice treated with CAR^+^ NKG2C^+^ NK-like T cells and CAR^+^ conventional CD8 T cells, measured 4 hours pre- and 24 hours post-CAR treatment (n=14 mice/group). (D) Serum levels of human cytokines from mice treated with CAR^+^ NKG2C^+^ NK-like T cells and CAR^+^ conventional CD8 T cells, measured 4 hours pre- and 24 hours post-CAR treatment (n=14 mice/group). (E) Serum levels of murine cytokines from mice treated with CAR^+^ NKG2C^+^ NK-like T cells and CAR^+^ conventional CD8 T cells, measured 4 hours pre- and 24 hours post-CAR treatment (n=14 mice/group). Data are presented as mean values +/- SEM. Statistical analysis was performed by one-way or two-way ANOVA.

### TCR-redirected NKG2C^+^ NK-like T cells demonstrate superior in vitro killing of NY-ESO-1^+^/HLA-A*02^+^ A375 melanoma cells

We next modified NKG2C^+^ NK-like T cells with an affinity-enhanced TCR construct targeting an NY-ESO-1 tumor antigen-derived peptide presented by HLA-A*02 (*Figure 5A-B*). Retroviral transduction of NKG2C^+^ NK-like T cells yielded high expression of NY-ESO-1 TCR^+^ cells, as assessed by Vß13.1 staining (53.62% ± 9.26%), superior to that of conventional CD8 T cells (18.08% ± 3.45%) (*Figure 5A-B*). Transduced cells were then co-incubated with the NY-ESO-1^+^/HLA-A*02^+^ A375 melanoma cell line and compared for induction of activating receptors and exhaustion markers. TCR-transduced NKG2C^+^ NK-like T cells expressed lower levels of 4-1BB than their conventional CD8 T cell counterparts and, similar to CAR-transduced NK-like T cells, did not upregulate the exhaustion markers PD-1, LAG-3 and TIM-3 (*Figure 5C*). TCR-transduced NKG2C^+^ NK-like T cells also demonstrated superior cytolysis of A375 target cells, compared with TCR-transduced conventional CD8 T cells, demonstrating efficient redirection of tumor cytolysis capacity (*Figure 5D*).

**Figure 5.**
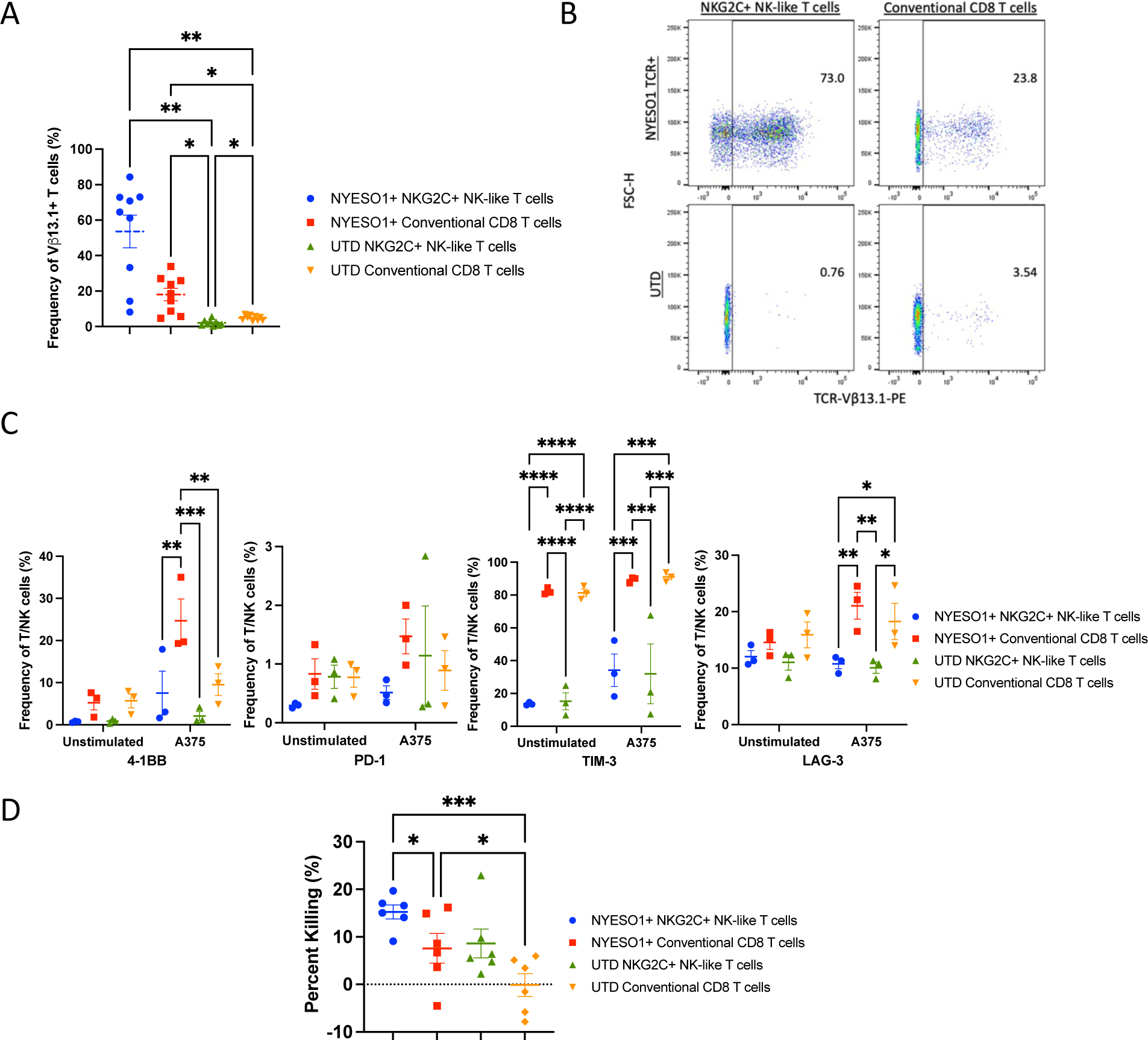
TCR-engineered NKG2C^+^ NK-like T cells mediate robust effector functions against NY-ESO-1^+^ melanoma target cells. (A) NY-ESO-1 TCR expression and (B) representative pseudocolor plots showing transduction efficiency measured via TCR Vß13.1 staining via flow cytometry (NY-ESO-1 TCR^+^ = Vß13.1^+^) (n=9 independent experiments). (C) Flow cytometry phenotype of TCR-engineered T cells with and without 8-hour stimulation with NY-ESO-1^+^ A375 target cells (n=3 independent experiments). (D) Cytotoxicity of TCR-engineered T cells against NYESO^+^ A375 target cells (n=6 independent experiments). Data are presented as mean values +/- SEM. Statistical analysis was performed by one-way or two-way ANOVA.

### TCR-transduced NKG2C^+^ NK-like T cells demonstrate in vivo tumor control of NY-ESO-1^+^/HLA-A*02^+^ A375 melanoma, comparable to conventional CD8 T cells

Immunodeficient NRG mice were then engrafted with A375 tumors via flank injection and were treated 10 days later with TCR-transduced NKG2C^+^ NK-like T cells, TCR-transduced conventional CD8 T or their untransduced counterparts (*Figure 6A*). Mice receiving TCR-transduced NKG2C^+^ NK-like T cells or TCR-transduced conventional CD8 T cells were comparable in slowing tumor growth, demonstrating both significant reduction in tumor volume and tumor weight at day 17 post-T cell treatment compared to control mice receiving untransduced T cells or tumor only (*Figure 6B-C*).

**Figure 6.**
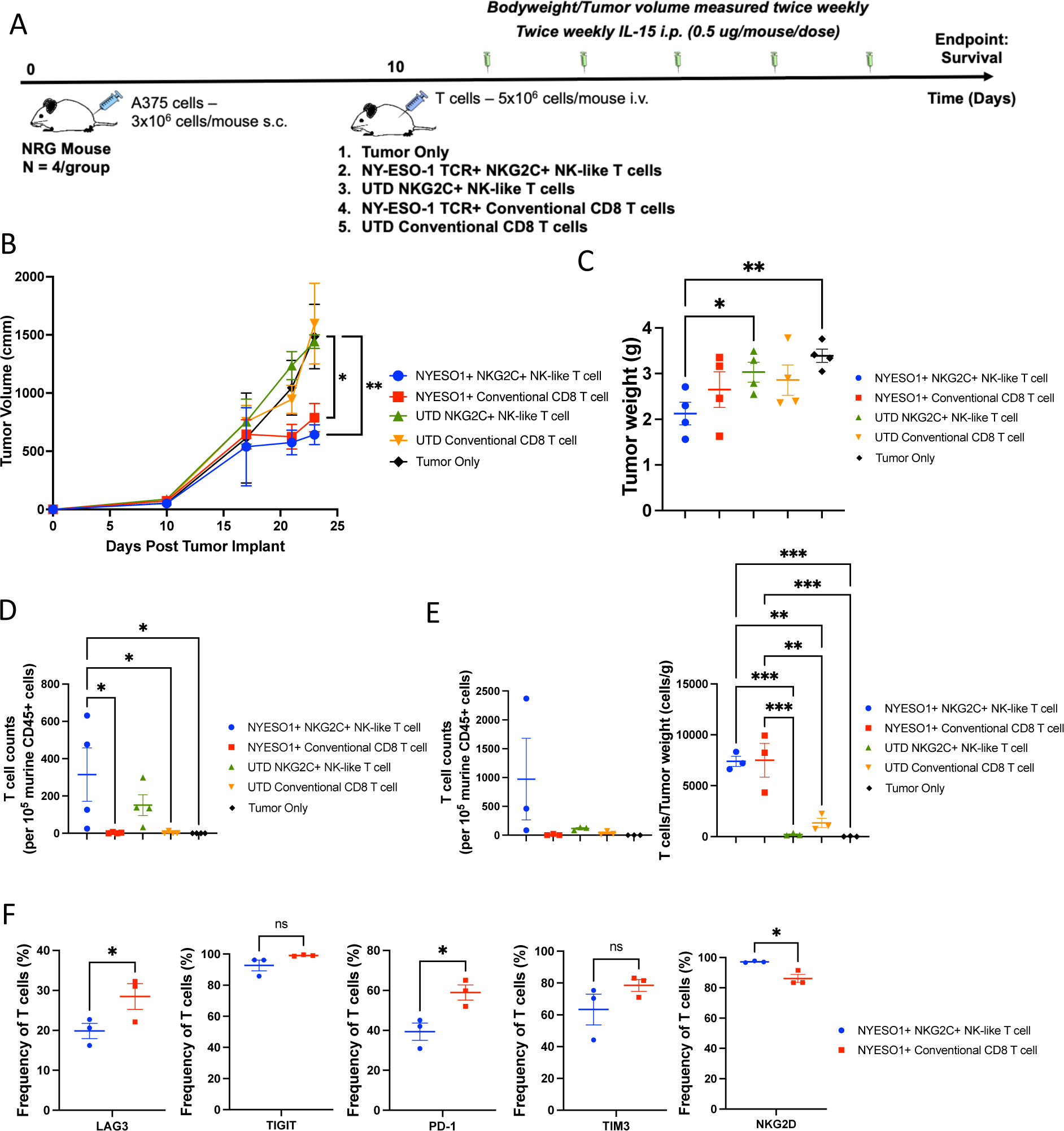
TCR-engineered NKG2C^+^ NK-like T cells mediate comparable tumor control and exhibit less exhaustion compared to TCR-engineered conventional CD8 T cells. (A) Study timeline of A375 tumor-bearing mice treated with NY-ESO-1 TCR-engineered or non-engineered NKG2C^+^ NK-like T cells and conventional CD8 T cells. (B) Tumor volume of mice treated with tumor only, TCR-engineered or non-engineered NKG2C^+^ NK-like T cells and conventional CD8 T cells, measured throughout the course of the study (n=4 mice/group). (C) Final tumor weight of mice treated with tumor only, TCR-engineered or non-engineered NKG2C^+^ NK-like T cells and conventional CD8 T cells, measured at day 17 post-treatment (n=4 mice/group). (D) Quantification of circulating TCR-engineered or non-engineered NKG2C^+^ NK-like T cells and conventional CD8 T cells at day 7 post-treatment (n=4 mice/group). (E) Quantification of circulating (left) and tumor infiltrating (right) TCR-engineered or non-engineered NKG2C^+^ NK-like T cells and conventional CD8 T cells at day 17 post-treatment (n=3 mice/group). (F) Frequency of activating and exhaustion markers of tumor infiltrating TCR-engineered or non-engineered NKG2C^+^ NK-like T cells and conventional CD8 T cells measured at day 17 post-treatment (n=3 mice/group). Data are presented as mean values +/- SEM. Statistical analysis was performed by one-way or two-way ANOVA or two-tailed unpaired t-test.

TCR-transduced NKG2C^+^ NK-like T cells exhibited higher persistence at day 7 and day 17 post-T cell infusion compared to TCR-transduced conventional CD8 T cells and untransduced control groups. While TCR-transduced NKG2C^+^ NK-like T cells had similar tumor infiltration to TCR-transduced conventional CD8 T cells at day 17 post-T cell infusion (*Figure 6D-E*), infiltrating TCR-transduced NKG2C^+^ NK-like T cells expressed lower levels of the exhaustion markers LAG-3 and PD-1 and higher levels of the activating receptor NKG2D (*Figure 6F, S16)*. These data demonstrate that while capable of comparable in vivo tumor control to conventional T cell therapies at a specified timepoint, NKG2C^+^ NK-like T cells are resistant to tumor-induced exhaustion.

## Discussion

NKG2C^+^ NK-like T cells represent a promising platform for engineered cell therapeutics, harnessing the advantageous characteristics of T and NK cell therapeutics such as durable in vivo persistence and innate tumor killing, while mitigating adverse effects such as CRS. We show that NKG2C^+^ NK-like T cells can be efficiently modified with a 1928z CAR construct or a transgenic TCR targeting the NY-ESO-1 antigen and rapidly expanded following retroviral transduction. CAR^+^ NKG2C^+^ NK-like T cells exhibit robust killing of CD19^+^ NALM6 and Raji cell lines in vitro and significantly slowed tumor progression in NALM6-bearing mice, resulting in improved survival over CAR^+^ conventional CD8 T cell- and CAR^+^ NK cell-treated mice. Further, CAR^+^ NKG2C^+^ NK-like T cells have superior in vivo persistence following transfer compared to CAR^+^ conventional CD8 T cells and CAR^+^ NK cells. This persistence is durable, even without the presence of CD4 T cells, a major contributor to CRS and a necessary requirement for in vivo persistence of conventional CD8 T cells^31, 32, 33^. Furthermore, NKG2C^+^ NK-like T cells, like NK cells, are sensitive and responsive to IL-15 treatment, which dramatically improves their in vivo persistence. Compellingly, these NK-like T cells do not upregulate exhaustion markers, such as PD-1, TIM-3 and LAG-3, suggesting that they are not only powerful effector cells with innate tumor targeting capabilities, but also that they are resistant to tumor-induced exhaustion mechanisms, distinguishing these cells as a superior platform for engineered cell therapeutics. Using an in vivo CRS model, we further demonstrated that, likely due to their NK-like identity, CAR^+^ NKG2C^+^ NK-like T cells do not induce CRS, a hallmark consequence of traditional CAR-T cell therapies. The elevated levels of hIL-2 in 1928z^+^ NKG2C^+^ NK-like T cell-treated mice may be caused by their in vitro culture on feeder cells expressing mbIL-21, which has been shown to enhance IL-2 secretion in CD8 T cells^34^, while the production of hIL-2 in 1928z^+^ conventional CD8 T cell-treated mice is likely directly related to the CRS response, consistent with the upregulation of other human and murine cytokines. Taken together, the results indicate that NKG2C^+^ NK-like T cells are not only a more cytolytic platform for cell engineering with greater potential for long-term persistence, but also a safer alternative to traditional cell therapeutics.

A recent Phase I clinical trial of 31 patients with large B cell lymphoma treated with a CD19-targeted CAR-T cell (ACTRN12617001579381) reported an expansion of NK-like CD8 T cells, which correlated with superior outcomes in treated patients, suggesting that, even in current T cell therapeutics, NK-like CD8 T cells may be a significant contributor to durable therapeutic outcomes^35^. Patients with complete response at 6 months had an enrichment for NK-like CD8 T cells, including those with transcripts for *KLRC2*, the gene encoding for NKG2C, compared to patients with progressive disease. These results further support the role of NK-like CD8 T cells in controlling disease and justify the use of an enriched NK-like CD8 T cell product in treating B cell and other hematologic malignancies and solid tumors and mitigating CRS.

TCR-transduced NKG2C^+^ NK-like T cells demonstrated superior in vitro effector functions against an A375 melanoma target cell line over TCR-transduced conventional CD8 T cells. In vivo, TCR-transduced NKG2C^+^ NK-like T cells demonstrated comparable tumor control and infiltration, with enhanced persistence and reduced expression of exhaustion markers. While TCR-engineered NKG2C^+^ NK-like T cells did not offer superior tumor control over similarly engineered conventional T cells at the specified endpoint, their resistance to exhaustion and durable in vivo persistence remain clear advantages over conventional T cell therapeutics. The superior in vitro function and superior exhaustion profile but comparable in vivo tumor control may be explained by reduced expression of CD28, the traditional TCR costimulatory receptor, on NKG2C^+^ NK-like T cells compared to conventional CD8 T cells^17^, a condition that can be circumvented through additional cell engineering strategies.

Overall, this study demonstrates that CAR- or TCR-modified NKG2C^+^ NK-like T cells are powerful effector cells, with NK-like innate effector function and cytokine response, and proliferation and persistence better than conventional CD8 T cells. Importantly, they display these advantages while exhibiting a superior safety profile to traditional T cell therapeutics. It should be noted that despite the population’s prominent expression of the NKG2C receptor, NKG2C is likely not responsible for the observed differences in safety, exhaustion, and cytotoxic profiles, as we have previously reported that NKG2C^+^ NK-like T cells maintain innate-like functions even after blocking NKG2C^17^. Instead, due to *BCL11B* downregulation, NKG2C^+^ NK-like T cells have a full innate-like phenotypic and transcriptional program, imparting these cells with the functional and safety characteristics of an NK cell^17^. From a manufacturing perspective, these cells are easily identifiable, isolated and expanded via their expression of the NKG2C molecule. They are efficiently modified via retroviral transduction and are easily expanded to sufficient numbers for adoptive cell therapeutics. While the question of whether these cells are superior as a cell therapy platform to other NK-like T cells, such as MAIT, γδ T, and KIR+ CD8 T cells^12, 14, 15, 18, 19, 21, 23, 36^, remains to be explored, NKG2C^+^ NK-like T cells offer a unique advantage by their natural expansion following CMV infection, a lifelong infection with frequent reactivation^17, 37^. Clonal expansion is mediated through both NKG2C as well as through the endogenous TCR, leading to a notably oligoclonal population, with one only one or two dominant clones in most individuals tested. The endogenous TCR exhibits HLA-E restriction in all evaluated donors and most likely recognizes a CMV-derived peptide^17^. Given the high global CMV seroprevalence, we propose that adoptively transferred modified NKG2C^+^ NK-like T cells will be maintained and potentially expanded in patients upon exposure to CMV, resulting in life-long persistence of these tumor-directed cells in patients^38^. Such enduring protection would not predictably occur with other non-CMV-driven NK-like T cells. Further, use of a K562-based, HLA-E-expressing feeder platform, exploits the NKG2C expression on this population leading to robust expansion of NKG2C^+^ NK-like T cells for the generation clinically relevant doses of modified T cells. These doses may be difficult to achieve with other NK-like CD8 T cell populations, which may be more rare and whose specificity is unknown. In patients who do not have naturally-occurring NKG2C^+^ NK-like T cells, innate NK-like T cells can be alternatively generated via deletion of *BCL11B*, resulting in a cell population with similar characteristics to NKG2C^+^ NK-like T cell population, thereby expanding the utility to a larger patient population^17, 27^.

In mouse models, persistence of xenografted NKG2C^+^ NK-like T cells was enhanced by their responsiveness to exogenous human IL-15. We hypothesize that upon transfer to patients, the anti-tumor efficacy we report in our study will be enhanced by readily available endogenous IL-15,^39, 40, 41^. Nevertheless, armoring of the CAR construct with rhIL-15 is also an acceptable strategy and currently employed for existing NK cell therapeutics to ensure durability and persistence following adoptive transfer^10, 28^. We have also previously reported that ex vivo expansion and cytolytic activity following deletion of the endogenous TCR remains robust and unimpeded, indicating that NKG2C^+^ NK-like T cells may be more amenable for allogeneic use than conventional T cell therapeutics^17^.

In vivo application of NKG2C^+^ NK-like T cells to target CD19 or NY-ESO-1 provides proof-of-concept of the ability to harness the robust effector functions of this promising cell type. Their innate-like target recognition, cytotoxicity, safety profile, and adaptive-like persistence in vivo support the use of NKG2C^+^ NK-like T cells as a durable platform for engineered cell strategies, accelerating meaningful clinical advancements in T cell therapeutics. Further, given the innate-like tumor recognition, enhanced proliferative capacity of these cells, and relative lack of exhaustion, we hypothesize that non-engineered NKG2C^+^ NK-like may also be useful in areas where non-engineered NK cells have proven promising, but remain short-lived. Currently, several clinical trials utilize cytokine-induced memory-like NK cells to treat acute myeloid leukemia (NCT06152809, NCT05580601, NCT06138587, NCT03068819, NCT02782546), multiple myeloma (NCT04634435), and head and neck squamous carcinoma (NCT04290546). NKG2C^+^ NK-like T cells offer a more persistent cell therapy option in these settings and ultimately may pave the way for more durable patient response.

## Methods

### Cell sources and preparation

Peripheral blood mononuclear cells (PBMCs) were isolated by Ficoll separation (GE Healthcare) from buffy coats obtained from New York Blood Center (NYBC). PBMCs were freshly isolated at the time of use or cryopreserved in fetal bovine serum (FBS) with 10% DMSO for later use. The MSKCC Institutional Review Board (IRB) has approved the use of these samples and has waived the need for additional research consent for anonymous NYBC samples. Human cytomegalovirus (HCMV) serostatus was provided by NYBC.

### Flow Cytometry, Cell Sorting, and Cell Isolation

Freshly isolated PBMCs were Fc-blocked (Miltenyi, 130-059-901) for 10 minutes and stained for the following surface markers for 30 minutes at room temperature: NKG2C-PE (R&D Systems, FAB138P-100; Miltenyi, 130-119-776), NKG2A-PE-Vio770 (Miltenyi, 130-113-567), CD3-BV650 (BD Biosciences, 563916), CD56-ECD (Beckman Coulter, A82943), CD8-BV786 (BD Biosciences, 563823), TCRαβ-APC (BD Biosciences, 563826), KIR2DL1/S1-FITC (Miltenyi, 130-118-973), KIR2DL2/L3/S2-FITC (BD Biosciences, 559784), KIR3DL1/S1-FITC (Miltenyi, 130-104-836). DAPI was added as a viability marker immediately prior to sorting. Donors with greater than 2% NKG2C^+^ NK-like T cells of all CD8 T cells were selected for evaluation, and cells were further purified for NKG2C^+^ and NKG2C^-^ cell populations. Briefly, PBMCs were Fc-blocked (Miltenyi, 130-059-901) for 10 minutes and stained for NKG2C-Biotin (Miltenyi, 130-120-448) for 30 minutes at room temperature. NKG2C^+^ cells were isolated using anti-Biotin UltraPure Microbeads (Miltenyi). The NKG2C^-^ fraction was subsequently isolated for CD8^+^ cells using anti-CD8 Microbeads (Miltenyi). Cells were allowed to rest overnight in RPMI supplemented with 10% FCS, 1% Penicillin-Streptomycin, and 200 IU/mL IL-2 (Peprotech). The following day, cells were FACS-sorted and two live cell populations were collected: NKG2C^+^ NK-like T cells (CD3^+^CD56^+^CD8^+^TCRαβ^+^NKG2C^+^NKG2A^-^) and conventional CD8 T cells (CD3^+^CD56^-^CD8^+^TCRαβ^+^NKG2C^-^NKG2A^-^KIR^-^) (*Figure S1*). For NK cells, cells were isolated by negative selection using the human NK Cell Isolation Kit (Miltenyi). Media for all cell lines was obtained from the MSKCC Media Preparation Core.

For subsequent immunophenotyping, isolated cells (2 x 10^5^ cells per well) were stained for 20 minutes with LIVE/DEAD^TM^ Fixable Aqua - Dead Cell Staining Kit (405 nm, Invitrogen) followed by 30 minutes staining of surface markers. Intracellular proteins were stained using the FIX & PERM^TM^ Cell Permeabilization Kit (Thermo Fisher). A full list of antibodies used is provided in supplementary *Table S1*.

### Cell Lines and culture

The human lymphoblastic leukemia cell line NALM6 (ATCC, CRL-3273) and human Burkitt lymphoma cell line Raji (ATCC, CCL-86) were cultured in RPMI supplemented with 10% FCS and 1% Penicillin-Streptomycin. The human erythroleukemia cell line K562 (ATCC, CCL-243) was sequentially transduced with HLA-E with the HLA-G*01 leader sequence peptide (VMAPRTLFL)^17^, membrane-bound IL-21 (mbIL-21)^42^, and 4-1BBL, following which it was single cell subcloned and cultured in RPMI supplemented with 10% FCS and 1% Penicillin-Streptomycin. The human erythroleukemia cell line K562 C9.mbIL-21, expressing mbIL-21, 4-1BBL, CD64, CD86, and truncated CD19^43^, was generously provided by Dr. Dean Lee (Nationwide Children’s Hospital, Columbus, OH), cultured in RPMI supplemented with 10% FCS and 1% Penicillin-Streptomycin, and used for expansion of NK cells. The human malignant melanoma cell line A375 (ATCC, CRL-1619) was cultured in DMEM supplemented with 10% FCS and 1% Penicillin-Streptomycin. NALM6, Raji, and A375 were stably transduced with a GFP-firefly luciferase fusion protein as described^44^. Further to generate CD19KO NALM6 target cells, NALM6 WT cells were transduced with a CRISPR-Cas9 lentiviral construct targeting CD19, pLV[CRISPR]-hCas9:T2A:Puro-U6>hCD19[gRNA#892] (Vector Builder), and selected for two weeks with 1 ug/mL puromycin. The HEK293GP cell line, expressing viral gal and pol proteins^45^, was cultured in DMEM supplemented with 10% FCS and 1% Penicillin-Streptomycin. The HEK293Galv9-1928z cell line, a stable retroviral producer line expressing the 1928z-mCherry construct^46^, was cultured in DMEM supplemented with 10% FCS and 1% Penicillin-Streptomycin. All cells were verified by morphology and were confirmed to be mycoplasma-free.

### Retroviral Production and Transduction

The 1928z CAR retroviral vector was generously provided by Dr. Renier Brentjens (Roswell Park Comprehensive Cancer Center, Buffalo, NY) and was generated using the SFG backbone, as previously described ^46, 47, 48^. The NY-ESO-1 transgenic TCR retroviral vector was generated using the MSGV backbone, as previously described ^45, 49, 50^. This vector consists of an optimized high-affinity 1G4 NY-ESO-1 directed TCR, as previously described ^45, 49, 50^.

Sorted NKG2C^+^ NK-like T cells were expanded with 1:1 co-culture with irradiated K562-HLA-E:G*01-mbIL21-41BBL in RPMI supplemented with 10% FCS, 1% Penicillin-Streptomycin, and 200 IU/mL IL-2 for one week prior to transduction. Sorted conventional CD8 T cells, were expanded 1:2 with CD3/2/28 beads (Miltenyi) in RPMI supplemented with 10% FCS, 1% Penicillin-Streptomycin, and 200 IU/mL IL-2 for one week prior to transduction. Isolated NK cells were expanded 1:1 with irradiated K562 C9.mbIL-21 in RPMI supplemented with 10% FCS, 1% Penicillin-Streptomycin, and 200 IU/mL IL-2 for one week prior to transduction. After 1 week of expansion, 10^6^ cells were spinoculated at 2000g for 1 hour on RetroNectin (Takara Clontech) coated plates with retroviral supernatant either from HEK293Galv9-1928z or from HEK293GP, transfected with the NY-ESO-1 TCR transgene and RD114 envelope vector. After 24 hours, this step was repeated, to generate CAR^+^ T and NK cells and TCR^+^ T cells. After 48 hours, cells were assessed for transduction efficiency, expanded, and used for assays. For expansion assays, cells were counted via trypan blue exclusion.

### Functional Assays

For cytolysis assays, engineered- and non-engineered-T and NK cells were cocultured with 1x10^4^ GFP-luciferase expressing target cells at various E:T ratios for 4 hours in triplicate in white-walled 96-well plates (Corning) in a total volume of 200 uL. Additional wells of targets alone were plated at the same density to determine the maximum luciferase expression. After co-incubation, 75 ng of D-Luciferin (Gold Biotechnology) dissolved in RPMI medium was added to each well and incubated for 10 minutes. Luminescence of each sample was recorded using a Spark plate reader (Tecan) and SparkControl software (Tecan). Percent lysis was determined by the following calculation: [(Target alone signal-sample signal)/(target alone signal)]x100.

For CD107a mobilization and IFN-γ production assays, 2x10^5^ engineered- and non-engineered-T and NK cells were cocultured 1:1 with target cells or alone in the presence of CD107a-BV786 antibody. After 1.5 hours of coculture, 55.5 ug/mL of Brefeldin A (MP Biomedicals) and 1uM of GolgiStop (BD) were added to cells. After an additional 4 hours, cells were washed, stained for cell surface markers, fixed, permeabilized, and stained for IFN-γ.

For immunophenotyping following target stimulation, 2x10^5^ engineered- and non-engineered- T and NK cells were cocultured at a 1:1 E:T ratio with or without target cells for 8 hours. Following coculture, cells were washed and stained for cell surface markers.

### Mice

Male and female 6- to 8-week-old NOD.Cg-Rag1^tm1Mom^IL2rg^tm1Wji^/SzJ (NRG) were purchased from The Jackson Laboratory and C.B.Tgh-1b/GbmsTac-Prkdc^scid^Lyst^bg^N7 (SCID-beige) mice were purchased from Charles River. Both mouse strains were maintained by the MSKCC mouse facility. All animal experiments described in this study were approved by the MSKCC Institutional Animal Care and Use Committee (IACUC), according to all relevant animal use guidelines and ethical regulations (#09-10-021). Mice were housed in a 12 hour/12 hour light/dark cycle, with a temperature of 18-23°C and relative humidity of 40-60%.

### In vivo xenograft models

For CAR-T efficacy studies, NALM6 cells (0.5 x 10^6^) were inoculated intravenously (i.v.) via tail vein injection into NRG mice. After 4 days, mice were randomly assigned to one of 13 experimental groups, with cells administered via tail vein injection: Tumor only control (n=5), 1928z^+^ NKG2C^+^ NK-like T cells (5 x 10^6^/mouse) with or without IL-15 (n=5 per group), 1928z^+^ conventional CD8 T cells with or without IL-15 (5 x 10^6^/mouse) (n=5 per group), 1928z^+^ NK cells with or without IL-15 (5 x 10^6^/mouse) (n=5 per group), untransduced (UTD) NKG2C^+^ NK-like T cells (5 x 10^6^/mouse) with or without IL-15 (n=5 per group), UTD conventional CD8 T cells with or without IL-15 (5 x 10^6^/mouse) (n=5 per group), and UTD NK cells with or without IL-15 (5 x 10^6^/mouse) (n=5 per group). Mice were treated with or without biweekly IL-15 co-administration (0.5 ug/mouse/injection) by intraperitoneal (i.p.) injection. Blood was collected via retro-orbital bleed at day 7 post-T/NK cell treatment, red blood cells were lysed, and cells were stained for activation and exhaustion markers and analyzed via flow cytometry. Mice were monitored for survival and bioluminescence throughout the study by an investigator blinded to individual treatment groups.

For in vivo CRS studies, Raji cells (3 x 10^6^) were inoculated via i.p. injection into SCID-Beige mice. After 21 days, mice were randomly assigned to one of three experimental groups, with effector cells administered via i.p. injection: Tumor only control (n=14), 1928z^+^ NKG2C^+^ NK-like T cells (30 x 10^6^/mouse) (n=14), and 1928z^+^ conventional CD8 T cells (30 x 10^6^/mouse) (n=14). Mice were monitored for 72 hours post-T cell injection for bodyweight, and blood serum was collected at 4 hours pre-treatment, and 24, 48, and 72 hours post treatment. Following 72 hours, mice were sacrificed, and blood, bone marrow, spleen, and peritoneal lavage samples were collected. Murine immune cells were isolated from blood, spleen, and tumors, and stained and analyzed via flow cytometry. Pre-treatment and 24 hour post-treatment serum was analyzed for human and serum cytokine levels using Cytokine Bead Array (BD) (hIL2, hGMCSF, hIL3, hIFNg, hIL6, hGCSF, hTNF, mIFNg, mIL3, mGMCSF, mIL5, mIL6, mCCL2, mGCSF, mCXCL9). Serum was also analyzed by ELISA for mCXCL1 (Thermo Fisher) and mSAA-3 (Millipore-Sigma).

For TCR-T efficacy studies, A375 melanoma cells (3 x 10^6^) were inoculated subcutaneously (s.c.) in the right flank of NRG mice. A375 melanoma cells (3 x 10^6^) we resuspended in 50 uL of PBS and mixed 1:1 by volume with 50 uL of Cultrex (Cultrex Basement Membrane Extract, Type 3, Pathclear, Bio-Techne) immediately prior to inoculation. Tumor dimensions (L, W, H) were monitored, and the tumor volume was calculated according to the formula: V = 0.52 × L × W × H. After 10 days, tumors reached a volume of about 80 mm^3^ and mice were then randomly assigned to one of five experimental groups, with effector cells administered via tail vein injection: Tumor only control (n=4), NY-ESO-1 TCR^+^ NKG2C^+^ NK-like T cells (5 x 10^6^/mouse) (n=4), NY-ESO-1 TCR^+^ conventional CD8 T cells (5 x 10^6^/mouse) (n=4), UTD NKG2C^+^ NK-like T cells (5 x 10^6^/mouse) (n=4), and UTD conventional CD8 T cells (5 x 10^6^/mouse) (n=4). Mice were treated with biweekly IL-15 co-administration (0.5 ug/mouse/injection) by i.p. injection. Mice were sacrificed at 17 days post-T cell treatment, and blood and tumors were harvested, stained for immunologic activation and exhaustion markers and analyzed via flow cytometry.

### Statistical Analysis

For comparisons of two groups, unpaired t-tests were applied. ANOVA with multiple comparisons were applied to analyze groups of more than two. All tests are indicated in each figure legend. *p <0.05, **p <0.01, ***p <0.001, and ****p <0.0001 were used as significant p values. Data was analyzed using Prism 9 software (GraphPad).

## Acknowledgments

The authors would like to acknowledge Dr. Jean-Benoît Le Ludeuc (MSKCC) for prior characterization of the NKG2C^+^ NK-like T cell evaluated throughout this study, and Dr. Michel Sadelain for helpful discussions.

## Funding

The following funding was used to support these studies: R01 AI150999 (K.B.L., M.K.P., S.S., R.S., C.L., G.K., T.K., K.C.H.), MSKCC Center for Experimental Therapeutics Big Bets program (K.B.L., M.K.P., S.S., R.S., C.L., G.K., T.K., K.C.H.), MSKCC Technology Development Fund (K.B.L., M.K.P., S.S., R.S., C.L., G.K., T.K., K.C.H.), the Major Family Fund (K.B.L., M.K.P., S.S., R.S., C.L., G.K., T.K., K.C.H.); NIH/NCI Cancer Center Support Grant P30 CA008748 (K.B.L., M.K.P., S.S., R.S., C.L., G.K., T.K., K.C.H.). C.A.K. was supported in part by National Institutes of Health (NIH) grants R37 CA259177, R01 CA269733, R01 CA286507, P50 CA217694, The Sarcoma Center at Memorial Sloan Kettering Cancer Center, Cycle for Survival, and the Parker Institute for Cancer Immunotherapy.

## Author contributions

K.B.L., M.K.P., R.S., and K.C.H. conceived, designed and executed the experiments and wrote the manuscript; K.B.L., C.L., G.K., and T.K. maintained mice and carried out *in vivo* xenograft studies; S.S. generated the K562-HLA-E-expressing cell line used for expansion; A.F.D., S.S.C., and C.A.K. provided key reagents and contributed to experimental design; all authors reviewed and edited the manuscript.

## Competing interests

K.B.L. is listed as an inventor on a patent for targeting TIGIT and CD73 using engineered NK cells (patent number US-20230407257-A1 from US Patent and Trademark Office). M.K.P., R.S., and K.C.H. are listed as inventors on a provisional patent for the manufacture and use of NKG2C^+^CD8^+^ T cells (US provisional patent application number: 63/188,435). M.K.P. is listed as an inventor on a patent for targeting CD20 in canine malignancies (patent number 11116795 from US Patent and Trademark Office). M.K.P and K.C.H. are listed as inventors on a provisional patent for the design and use of HLA-E:peptide chimeric molecules used in the K562 expansion platform (patent number US-20240200031-A1 from US Patent and Trademark Office). K.C.H. is an the scientific advisor board for Wugen, Inc. and is a consultant for Exelixis. A.F.D. is a co-inventor of intellectual property related to CD371 CAR T-cell technology and field of use specific for allogeneic cell therapies, licensed by MSK to Caribou Biosciences. C.A.K. and S.S.C. are inventors on patents related to T cell receptor (TCR) discovery and public neoantigen-specific TCRs and are recipients of licensing revenue from Intima Bioscience shared according to Memorial Sloan Kettering Cancer Center (MSKCC) institutional policies. C.A.K. has consulted for or is on the scientific advisory boards for Achilles Therapeutics, Affini-T Therapeutics, Aleta BioTherapeutics, Bellicum Pharmaceuticals, Bristol Myers Squibb, Catamaran Bio, Cell Design Labs, Decheng Capital, G1 Therapeutics, Klus Pharma, Obsidian Therapeutics, PACT Pharma, Roche/Genentech and T-knife, and is a scientific co-founder and equity holder in Affini-T Therapeutics. S.S.C. is a scientific advisor and equity holder in Affini-T Therapeutics.

## Data and materials availability

Source data for all figures are provided with this paper and are available online. All other data supporting the findings are available from the corresponding author upon reasonable request.

## Notes

### Summary of Updates

Manuscript formatting updated to correct missing statistical comparisons in initial submission

## References

1. Yin, X., He, L. & Guo, Z. T-cell exhaustion in CAR-T-cell therapy and strategies to overcome it. Immunology 169, 400–411 (2023).

2. Teoh, P.J. & Chng, W.J. CAR T-cell therapy in multiple myeloma: more room for improvement. Blood Cancer J 11, 84 (2021).

3. Messmer, A.S. et al. CAR T-cell therapy and critical care : A survival guide for medical emergency teams. Wien Klin Wochenschr 133, 1318–1325 (2021).

4. Cai, C. et al. A comprehensive analysis of the fatal toxic effects associated with CD19 CAR-T cell therapy. Aging (Albany NY) 12, 18741–18753 (2020).

5. First CAR-T therapy to target BCMA gets FDA nod. Nat Biotechnol 39, 531 (2021).

6. Mailapalli, S., Kumar, R., Suresh, P., Singhal, A. & Singh, S. Clinicopathological Profile and 2 Year Relapse Rates of Non-Hodgkin’s Lymphoma in Tertiary Care Center. J Assoc Physicians India 70, 11–12 (2022).

7. Harrysson, S. et al. Incidence of relapsed/refractory diffuse large B-cell lymphoma (DLBCL) including CNS relapse in a population-based cohort of 4243 patients in Sweden. Blood Cancer J 11, 9 (2021).

8. Montserrat, E. & Dreger, P. Hope for high-risk chronic lymphocytic leukemia relapsing after allogeneic stem-cell transplantation. J Clin Oncol 33, 1527–1529 (2015).

9. Pfefferle, A. & Huntington, N.D. You Have Got a Fast CAR: Chimeric Antigen Receptor NK Cells in Cancer Therapy. Cancers (Basel) 12 (2020).

10. Marin, D. et al. Safety, efficacy and determinants of response of allogeneic CD19-specific CAR-NK cells in CD19(+) B cell tumors: a phase 1/2 trial. Nat Med (2024).

11. Merino, A., Maakaron, J. & Bachanova, V. Advances in NK cell therapy for hematologic malignancies: NK source, persistence and tumor targeting. Blood Rev 60, 101073 (2023).

12. Bjorkstrom, N.K. et al. CD8 T cells express randomly selected KIRs with distinct specificities compared with NK cells. Blood 120, 3455–3465 (2012).

13. Guma, M. et al. The CD94/NKG2C killer lectin-like receptor constitutes an alternative activation pathway for a subset of CD8+ T cells. Eur J Immunol 35, 2071–2080 (2005).

14. Correia, M.P. et al. Distinct human circulating NKp30(+)FcepsilonRIgamma(+)CD8(+) T cell population exhibiting high natural killer-like antitumor potential. Proc Natl Acad Sci U S A 115, E5980–E5989 (2018).

15. Balin, S.J. et al. Human antimicrobial cytotoxic T lymphocytes, defined by NK receptors and antimicrobial proteins, kill intracellular bacteria. Sci Immunol 3 (2018).

16. Mingari, M.C. et al. Human CD8+ T lymphocyte subsets that express HLA class I-specific inhibitory receptors represent oligoclonally or monoclonally expanded cell populations. Proc Natl Acad Sci U S A 93, 12433–12438 (1996).

17. Sottile, R., et al. Human cytomegalovirus expands a CD8(+) T cell population with loss of BCL11B expression and gain of NK cell identity. Sci Immunol 6, eabe6968 (2021).

18. Choi, S.J. et al. KIR(+)CD8(+) and NKG2A(+)CD8(+) T cells are distinct innate-like populations in humans. Cell Rep 42, 112236 (2023).

19. Li, J., et al. KIR(+)CD8(+) T cells suppress pathogenic T cells and are active in autoimmune diseases and COVID-19. Science 376, eabi9591 (2022).

20. Sullivan, L.C. et al. Natural killer cell receptors regulate responses of HLA-E-restricted T cells. Sci Immunol 6 (2021).

21. Good, C.R. et al. An NK-like CAR T cell transition in CAR T cell dysfunction. Cell 184, 6081–6100 e6026 (2021).

22. Giles, J.R. et al. Shared and distinct biological circuits in effector, memory and exhausted CD8(+) T cells revealed by temporal single-cell transcriptomics and epigenetics. Nat Immunol 23, 1600–1613 (2022).

23. Billiet, L. et al. Single-cell profiling identifies a novel human polyclonal unconventional T cell lineage. J Exp Med 220 (2023).

24. Ikawa, T. et al. An essential developmental checkpoint for production of the T cell lineage. Science 329, 93–96 (2010).

25. Kojo, S. et al. Priming of lineage-specifying genes by Bcl11b is required for lineage choice in post-selection thymocytes. Nat Commun 8, 702 (2017).

26. Li, L., Leid, M. & Rothenberg, E.V. An early T cell lineage commitment checkpoint dependent on the transcription factor Bcl11b. Science 329, 89–93 (2010).

27. Li, P. et al. Reprogramming of T cells to natural killer-like cells upon Bcl11b deletion. Science 329, 85–89 (2010).

28. Liu, E. et al. Cord blood NK cells engineered to express IL-15 and a CD19-targeted CAR show long-term persistence and potent antitumor activity. Leukemia 32, 520–531 (2018).

29. Giavridis, T. et al. CAR T cell-induced cytokine release syndrome is mediated by macrophages and abated by IL-1 blockade. Nat Med 24, 731–738 (2018).

30. Davila, M.L. et al. Efficacy and toxicity management of 19-28z CAR T cell therapy in B cell acute lymphoblastic leukemia. Sci Transl Med 6, 224ra225 (2014).

31. Turtle, C.J. et al. CD19 CAR-T cells of defined CD4+:CD8+ composition in adult B cell ALL patients. J Clin Invest 126, 2123–2138 (2016).

32. Melenhorst, J.J. et al. Decade-long leukaemia remissions with persistence of CD4(+) CAR T cells. Nature 602, 503–509 (2022).

33. Bove, C., et al. CD4 CAR-T cells targeting CD19 play a key role in exacerbating cytokine release syndrome, while maintaining long-term responses. J Immunother Cancer 11 (2023).

34. Yi, J.S., Ingram, J.T. & Zajac, A.J. IL-21 deficiency influences CD8 T cell quality and recall responses following an acute viral infection. J Immunol 185, 4835–4845 (2010).

35. Louie, R.H.Y. et al. CAR(+) and CAR(-) T cells share a differentiation trajectory into an NK-like subset after CD19 CAR T cell infusion in patients with B cell malignancies. Nat Commun 14, 7767 (2023).

36. Wang, C.Q., Lim, P.Y. & Tan, A.H. Gamma/delta T cells as cellular vehicles for anti-tumor immunity. Front Immunol 14, 1282758 (2023).

37. Sinclair, J. & Sissons, P. Latency and reactivation of human cytomegalovirus. J Gen Virol 87, 1763–1779 (2006).

38. Zuhair, M. et al. Estimation of the worldwide seroprevalence of cytomegalovirus: A systematic review and meta-analysis. Rev Med Virol 29, e2034 (2019).

39. Carson, W.E. et al. Endogenous production of interleukin 15 by activated human monocytes is critical for optimal production of interferon-gamma by natural killer cells in vitro. J Clin Invest 96, 2578–2582 (1995).

40. Silvestre, R.N. et al. Engineering NK-CAR.19 cells with the IL-15/IL-15Ralpha complex improved proliferation and anti-tumor effect in vivo. Front Immunol 14, 1226518 (2023).

41. Kochenderfer, J.N. et al. Lymphoma Remissions Caused by Anti-CD19 Chimeric Antigen Receptor T Cells Are Associated With High Serum Interleukin-15 Levels. J Clin Oncol 35, 1803–1813 (2017).

42. Singh, H. et al. Reprogramming CD19-specific T cells with IL-21 signaling can improve adoptive immunotherapy of B-lineage malignancies. Cancer Res 71, 3516–3527 (2011).

43. Denman, C.J. et al. Membrane-bound IL-21 promotes sustained ex vivo proliferation of human natural killer cells. PLoS One 7, e30264 (2012).

44. Brennan, T.V., Lin, L., Huang, X. & Yang, Y. Generation of Luciferase-expressing Tumor Cell Lines. Bio Protoc 8 (2018).

45. Wargo, J.A. et al. Recognition of NY-ESO-1+ tumor cells by engineered lymphocytes is enhanced by improved vector design and epigenetic modulation of tumor antigen expression. Cancer Immunol Immunother 58, 383–394 (2009).

46. Pegram, H.J. et al. IL-12-secreting CD19-targeted cord blood-derived T cells for the immunotherapy of B-cell acute lymphoblastic leukemia. Leukemia 29, 415–422 (2015).

47. Brentjens, R.J. et al. Eradication of systemic B-cell tumors by genetically targeted human T lymphocytes co-stimulated by CD80 and interleukin-15. Nat Med 9, 279–286 (2003).

48. Smith, E.L. et al. GPRC5D is a target for the immunotherapy of multiple myeloma with rationally designed CAR T cells. Sci Transl Med 11 (2019).

49. Cohen, C.J., Zhao, Y., Zheng, Z., Rosenberg, S.A. & Morgan, R.A. Enhanced antitumor activity of murine-human hybrid T-cell receptor (TCR) in human lymphocytes is associated with improved pairing and TCR/CD3 stability. Cancer Res 66, 8878–8886 (2006).

50. Cohen, C.J. et al. Enhanced antitumor activity of T cells engineered to express T-cell receptors with a second disulfide bond. Cancer Res 67, 3898–3903 (2007).

