## Supplemental Figures and Tables for "Engineered NKG2C^+^ NK-like T cells exhibit superior antitumor efficacy while mitigating cytokine release syndrome"

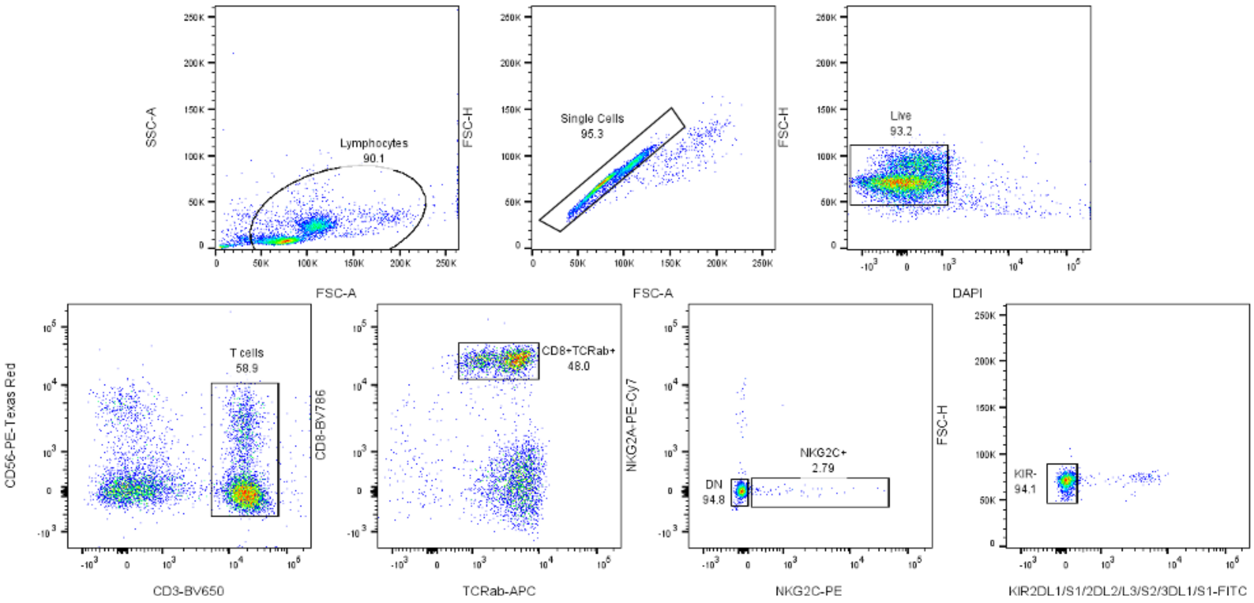


Figure S1. Flow cytometry gating strategy for FACS-sorted NKG2C⁺ NK-like T cells and conventional CD8 T cells.

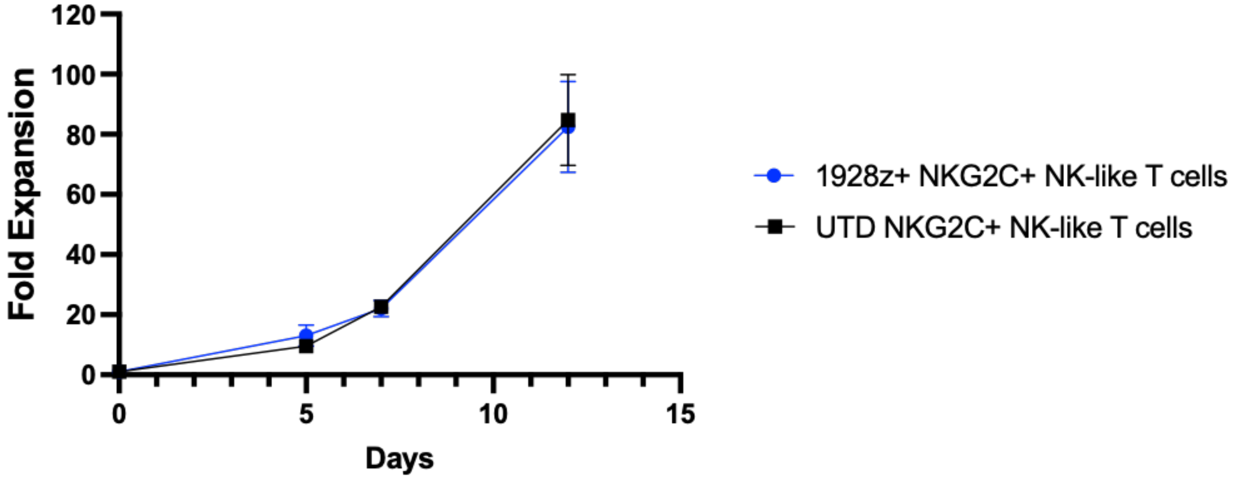


Figure S2. Fold ex vivo expansion of 1928z$^{+}$ NKG2C$^{+}$ NK-like T cells and untransduced (UTD) NKG2C$^{+}$ NK-like T cells. Data are presented as mean values +/- SEM. Statistical analysis was performed by multiple two-tailed unpaired t tests (n=3 independent experiments).

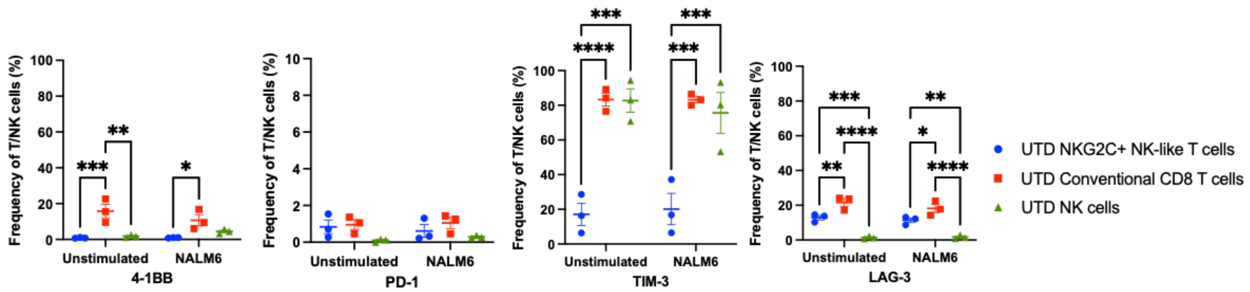


Figure S3. Flow cytometry phenotyping of UTD T and NK cells with or without 8-hour stimulation with $CD19^{+}$ NALM6 target cells. Data are presented as mean values +/- SEM. Statistical analysis was performed by two-way ANOVA (n=3 independent experiments).

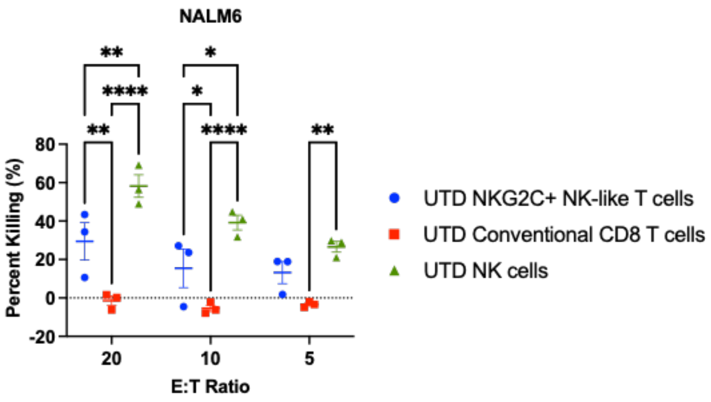


Figure S4. Cytotoxicity of UTD T and NK cells against $CD19^{+}$ NALM6 target cells. Data are presented as mean values +/- SEM. Statistical analysis was performed by two-way ANOVA (n=3 independent experiments).

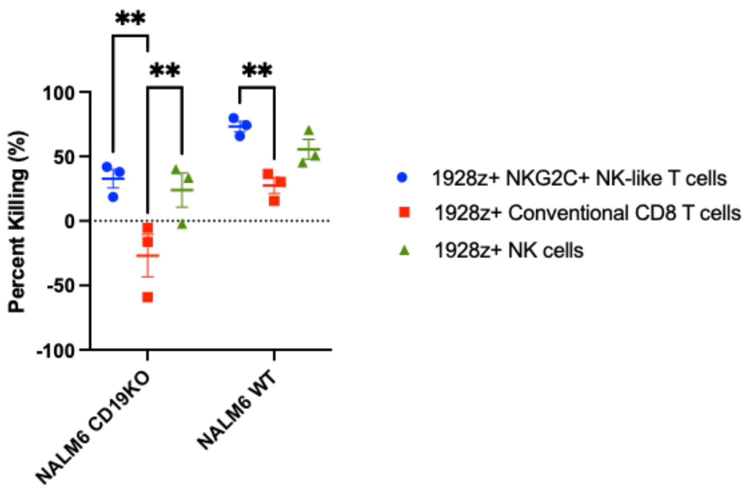


Figure S5. Cytotoxicity of 1928z$^{+}$ T and NK cells against CD19$^{+}$ NALM6 WT and NALM6 CD19KO target cells at a 5:1 E:T ratio. Data are presented as mean values +/- SEM. Statistical analysis was performed by two-way ANOVA (n=3 independent experiments).

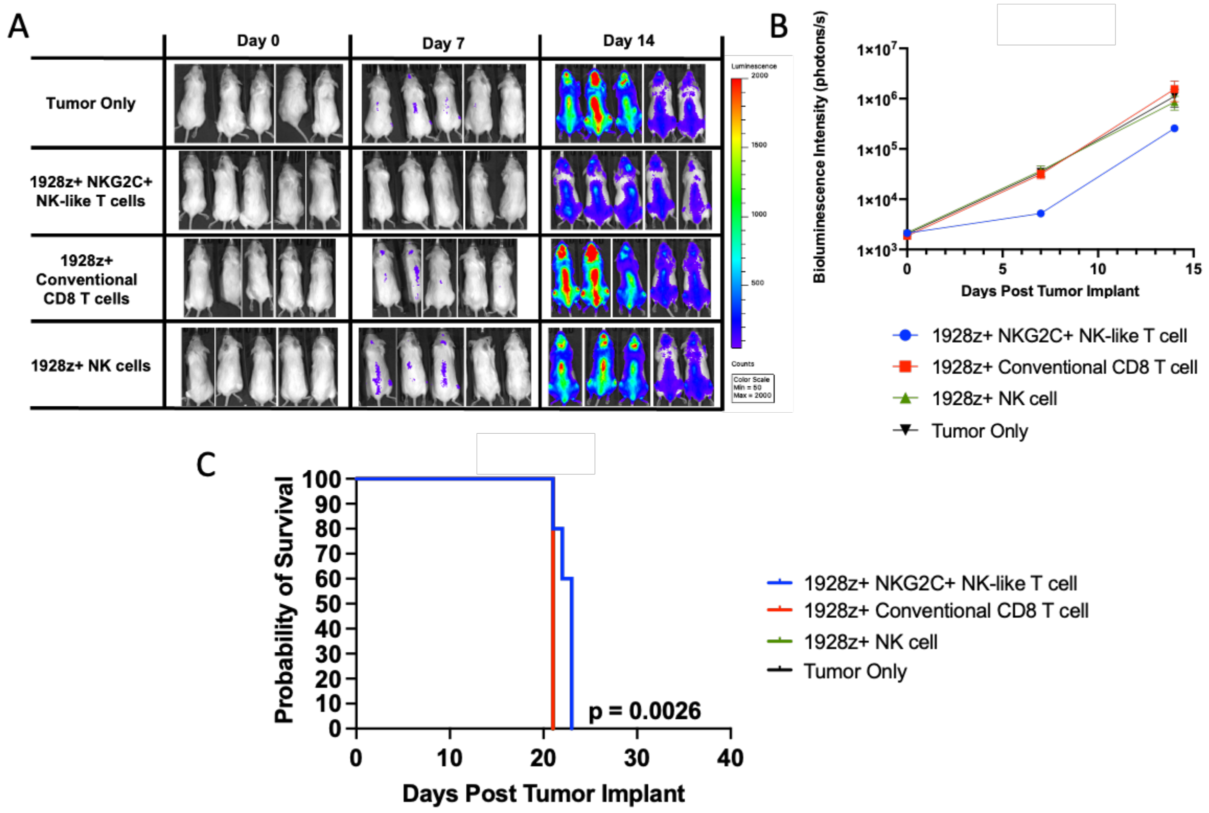


Figure S6. (A) Bioluminescence imaging of mice treated with tumor only, 1928z$^+$ NKG2C$^+$ NK-like T cells, 1928z$^+$ conventional CD8 T cells, and 1928z$^+$ NK cells without IL-15, measured throughout the course of the study. (B) Quantification of bioluminescence imaging of mice treated with tumor only, 1928z$^+$ NKG2C$^+$ NK-like T cells, 1928z$^+$ conventional CD8 T cells, and 1928z$^+$ NK cells, measured throughout the course of the study. (C) Kaplan-Meier survival plot of mice treated with tumor only, 1928z$^+$ NKG2C$^+$ NK-like T cells, 1928z$^+$ conventional CD8 T cells, and 1928z$^+$ NK cells, measured throughout the course of the study. Data are presented as mean values +/- SEM. Statistical analysis was performed by two-way ANOVA or Kaplan-meier survival analysis (n=5 mice/group).

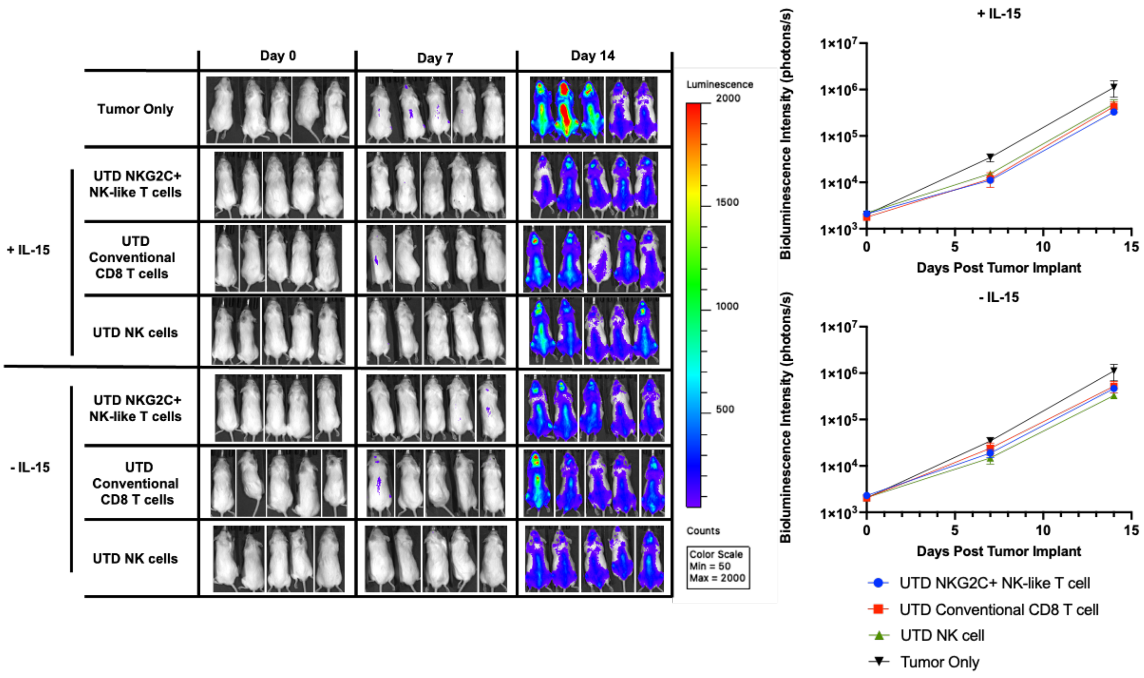


Figure S7. Bioluminescence imaging of mice treated with tumor only, UTD NKG2C$^{+}$ NK-like T cells, UTD conventional CD8 T cells, and UTD NK cells, measured throughout the course of the study (left). Quantification of bioluminescence imaging of mice, measured throughout the course of the study (right). Data are presented as mean values +/- SEM. Statistical analysis was performed by two-way ANOVA (n=5 mice/group).

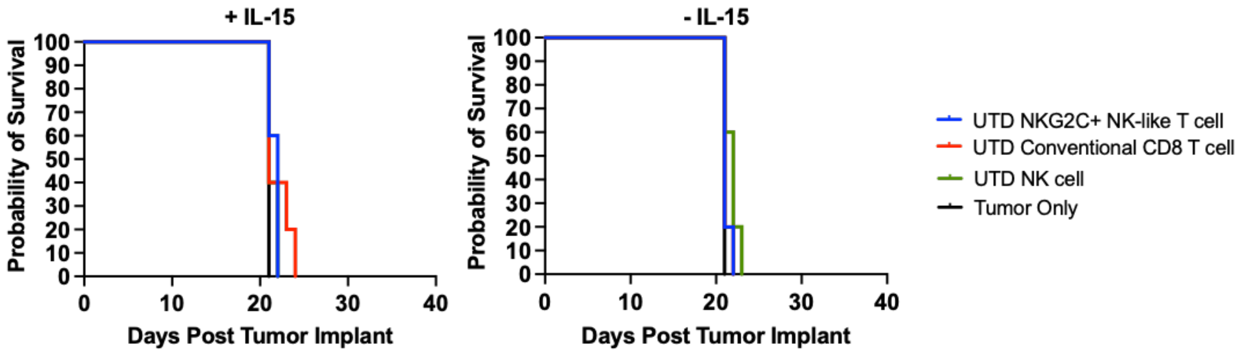


Figure S8. Kaplan-Meier survival plot of tumor only, and untransduced (UTD) T and NK cell groups treated with or without IL-15 co-administration. Statistical analysis was performed by Kaplan-Meier survival analysis (n=5 mice/group).

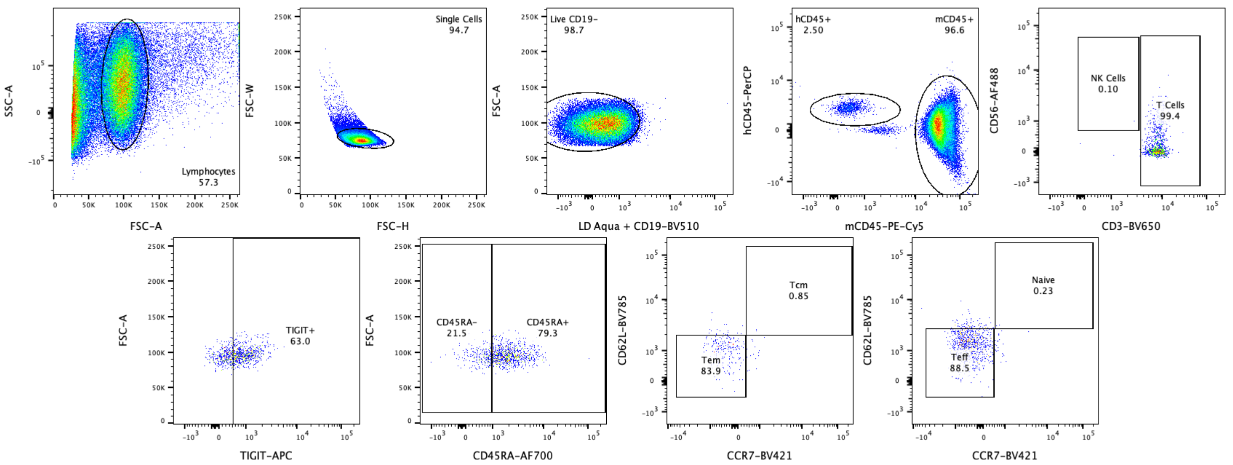


Figure S9. Flow cytometry gating strategy of T and NK cells collected from NALM6 or A375 tumor-bearing mice.

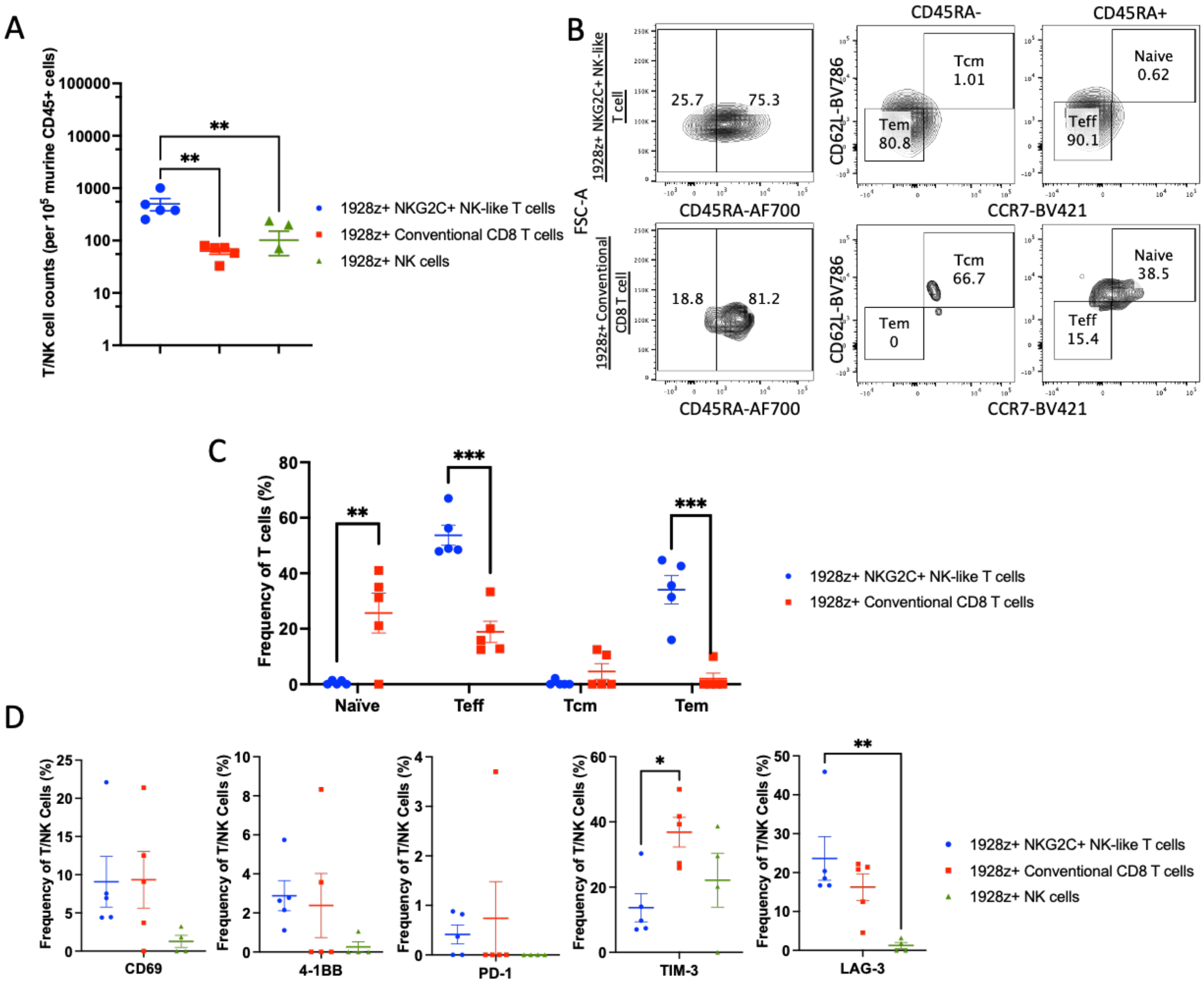


Figure S10. (A) Quantification of circulating CAR$^+$ NKG2C$^+$ NK-like T cells, CAR$^+$ conventional CD8 T cells, and CAR$^+$ NK cells without IL-15 at day 7 post-infusion, measured via flow cytometry. (B) Representative contour plots of circulating naïve, Teff, Tem, and Tcm populations among CAR$^+$ T cells at day 7 post-infusion, measured via CD45RA/CCR7/CD62L flow cytometry staining (C) Quantification of circulating naïve, Teff, Tem, and Tcm populations among CAR$^+$ T cells at day 7 post-infusion, measured via CD45RA/CCR7/CD62L flow cytometry staining (D) Activation and exhaustion marker phenotype of circulating CAR$^+$ T and NK cells at day 7 post-infusion, measured via flow cytometry. Data are presented as mean values +/- SEM. Statistical analysis was performed by one-way ANOVA or two-tailed unpaired t-test (n=5 mice/group).

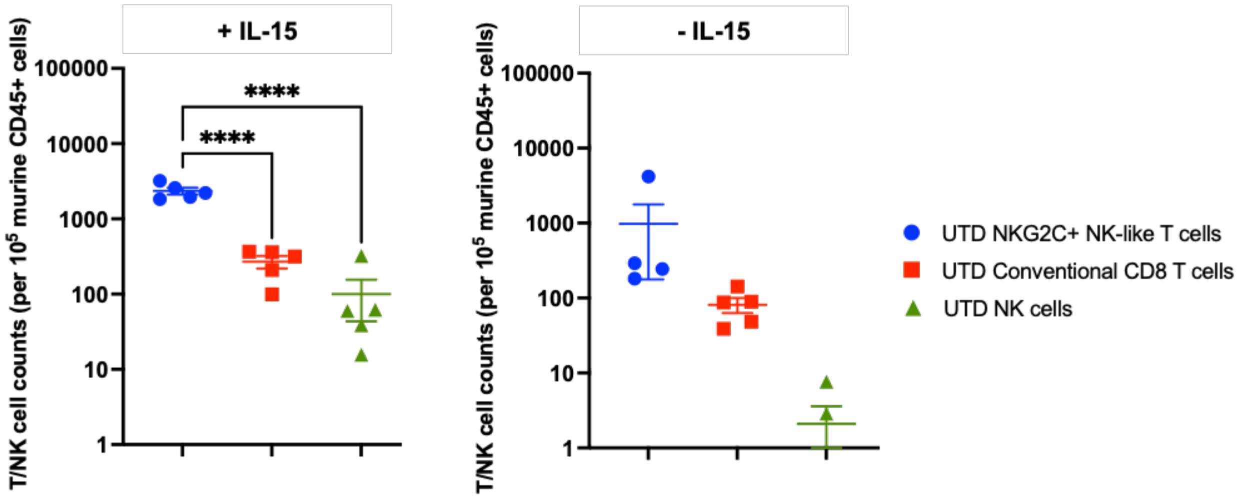


Figure S11. Quantification of circulating UTD NKG2C$^{+}$ NK-like T cells, UTD conventional CD8 T cells, and UTD NK cells at day 7 post-infusion, measured via flow cytometry. Data are presented as mean values +/- SEM. Statistical analysis was performed by one-way ANOVA (n=5 mice/group).

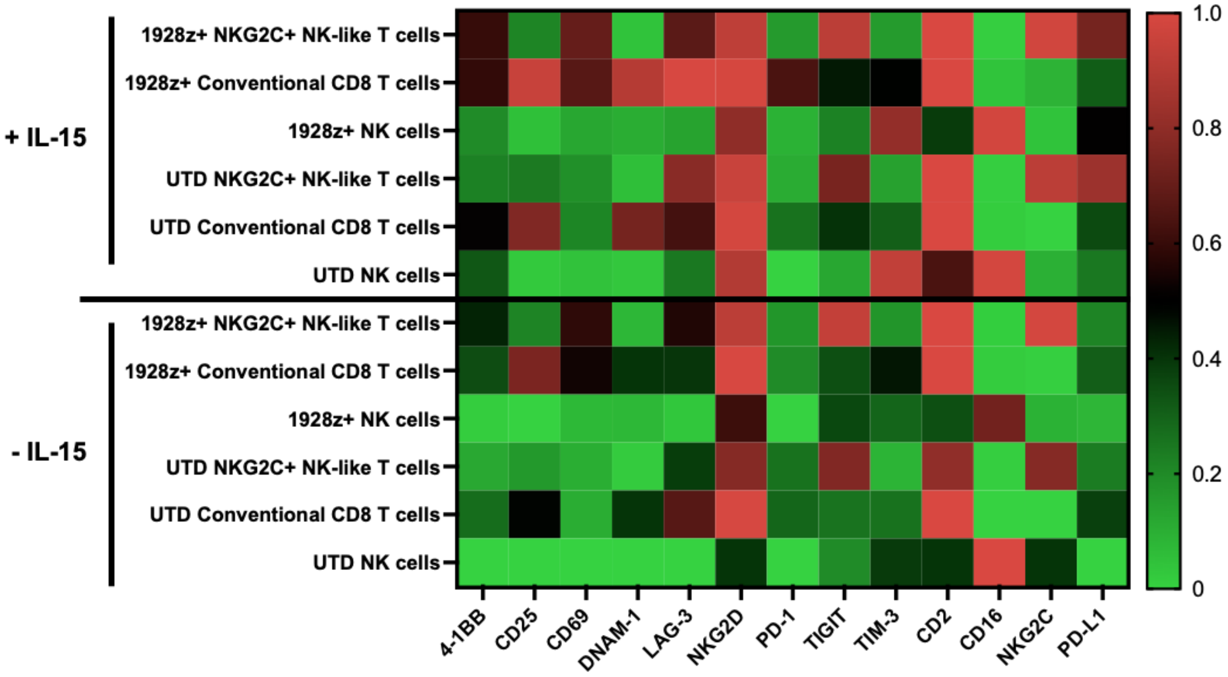


Figure S12. Heatmap of relative frequency of surface marker expression measured on T and NK cells collected from NALM6 tumor-bearing mice at day 7 post-infusion (n=5 mice/group).

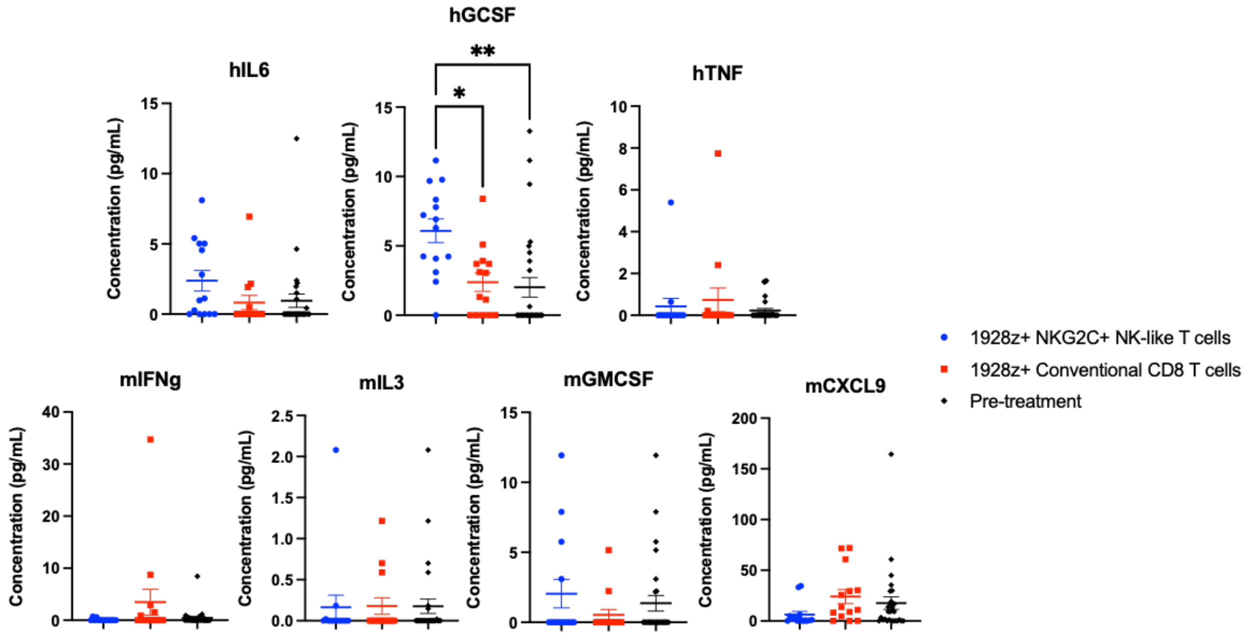


Figure S13. Serum levels of human and murine cytokines of mice treated with CAR$^+$ NKG2C$^+$ NK-like T cells and CAR$^+$ conventional CD8 T cells, measured at 4 hours pre- and 24 hours post-CAR infusion. Data are presented as mean values +/- SEM. Statistical analysis was performed by one-way ANOVA (n=14 mice/group).

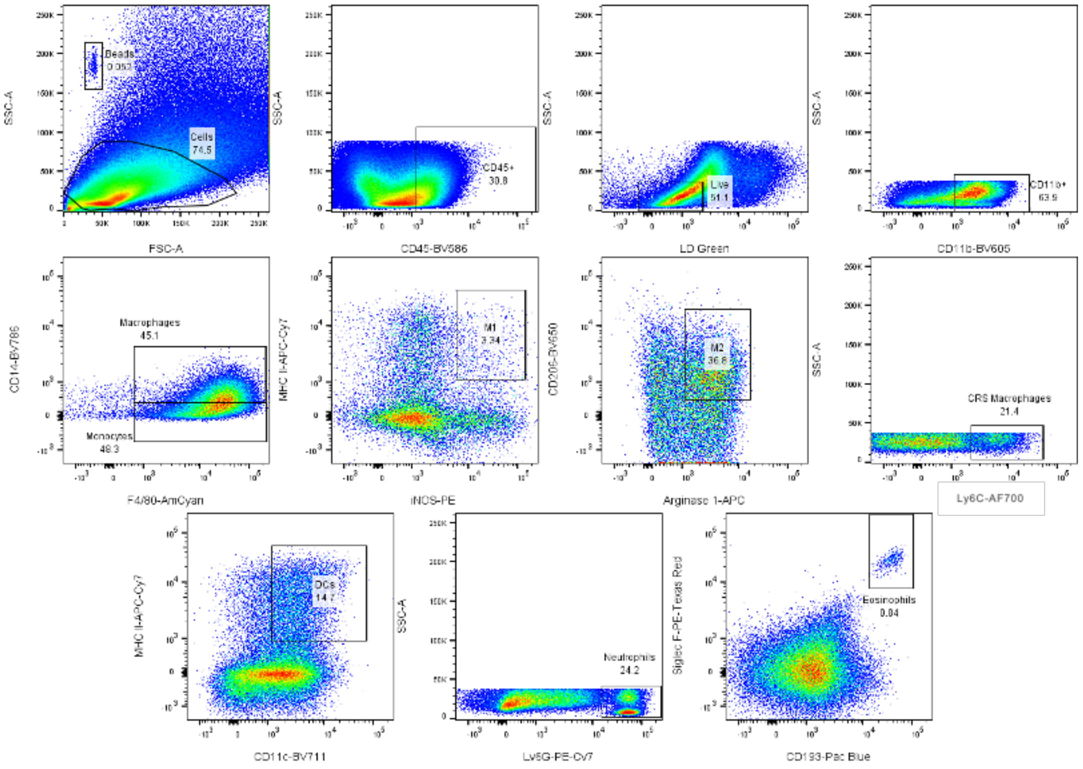


Figure S14. Flow cytometry gating strategy for murine myeloid subsets measured in bone marrow, blood, spleen, and peritoneal cavity of Raji tumor-bearing mice.

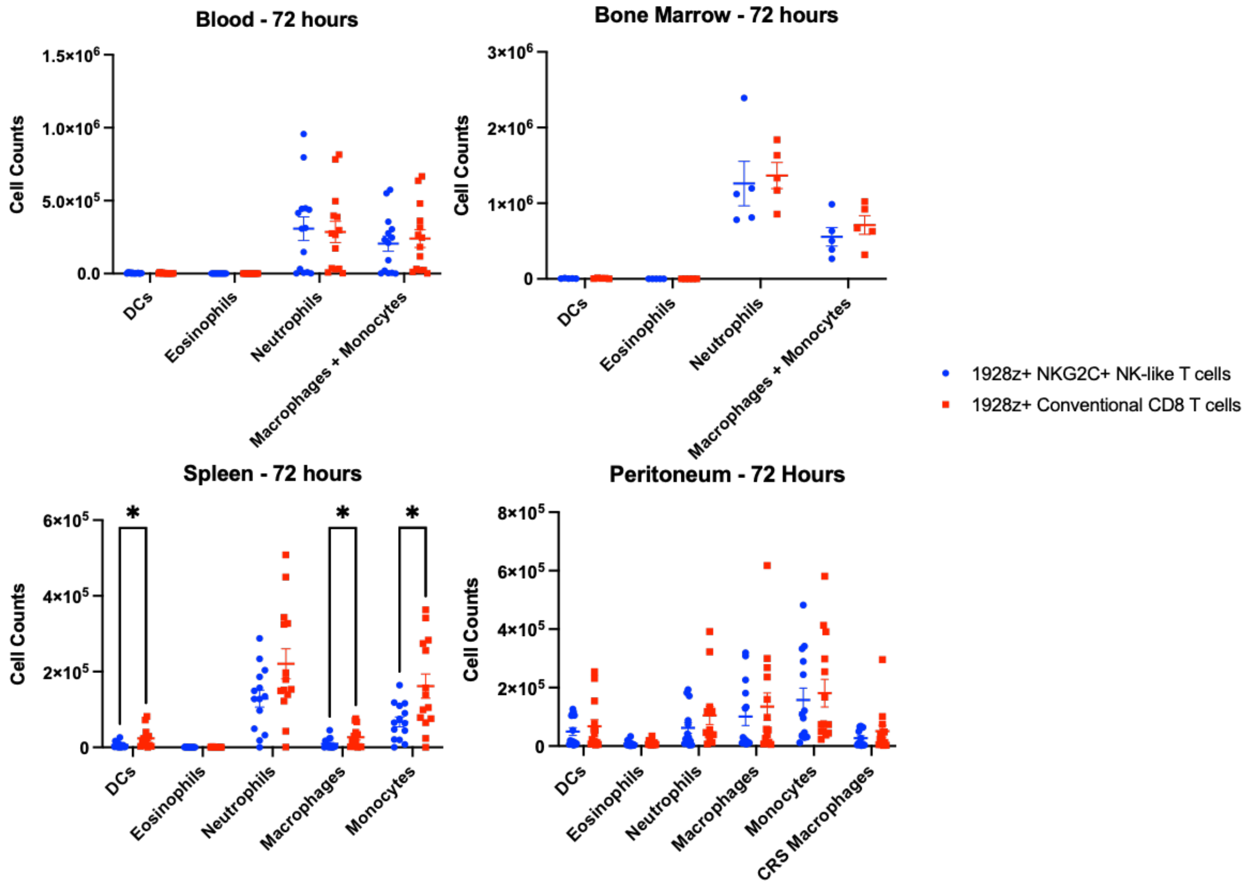


Figure S15. Quantification of murine myeloid subsets measured in bone marrow (BM), blood, spleen, and peritoneal cavity of Raji tumor-bearing mice 72 hours post T cell infusion. CRS macrophages are defined by expression of Ly6C. Data are presented as mean values +/- SEM. Statistical analysis was performed by multiple two-tailed unpaired t-tests (n=14 mice/group).

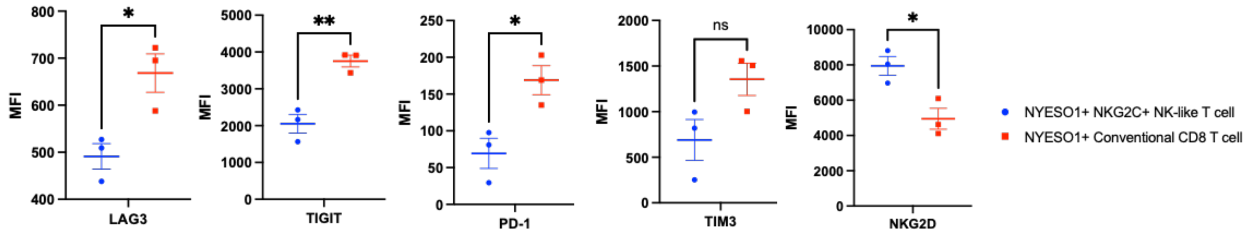


Figure S16. Median fluorescence intensity (MFI) of activating and exhaustion markers on tumor infiltrating TCR-engineered NKG2C$^{+}$ NK-like T cells and conventional CD8 T cells, measured at day 17 post-infusion. Data are presented as mean values +/- SEM. Statistical analysis was performed by two-tailed unpaired t-test (n=3 mice/group).

Table S1. Flow cytometry phenotyping antibodies and reagents.

| Antibody Marker/Reagent | Fluorescent Conjugate | Clone | Catalog Number | Manufacturer |
|---|---|---|---|---|
| TIGIT | AF647 | A15153G | 372724 | Biolegend |
| DNAM-1 | AF700 | 102511 | FAB666N-100UG | R&D Systems |
| PD-1 | APC-Fire750 | EH12.2H7 | 329954 | Biolegend |
| NKG2D | BV421 | 1D11 | 568106 | BD Biosciences |
| hCD19 | BV510 | SJ25C1 | 363020 | Biolegend |
| TIM-3 | BV605 | F38-2E2 | 345017 | Biolegend |
| CD3 | BV650 | UCHT1 | 563852 | BD Biosciences |
| LAG-3 | BV711 | 11C3C65 | 369320 | Biolegend |
| CD25 | BV785 | M-A251 | 356140 | Biolegend |
| CD69 | PE | FN50 | 310906 | Biolegend |
| mCD45 | PE-Cy5 | 30-F11 | 103110 | Biolegend |
| 4-1BB | PE-Cy7 | 4B4-1 | 309818 | Biolegend |
| CD56 | AF488 | B159 | 557699 | BD Biosciences |
| hCD45 | PerCP | 2D1 | 347464 | BD Biosciences |
| NKG2C | APC | REA205 | 130-117-398 | Miltenyi |
| CD45RA | AF700 | HI100 | 304120 | Biolegend |
| CD16 | APC-Cy7 | 3G8 | 557758 | BD Biosciences |
| CCR7 | BV421 | G043H7 | 353208 | Biolegend |
| CD2 | BV711 | RPA-2.10 | 300232 | Biolegend |
| CD62L | BV785 | DREG-56 | 304830 | Biolegend |
| PD-L1 | PE-Cy7 | 29E.2A3 | 329718 | Biolegend |
| TCR Vb13.1 | PE | REA560 | 130-131-910 | Miltenyi |
| CD107a | BV786 | H4A3 | 563869 | BD Biosciences |
| IFNy | AF700 | B27 | 557995 | BD Biosciences |
| CD56 | ECD | N901 | A82943 | Beckman Coulter |
| TCRab | APC | IP26 | 306718 | Biolegend |
| CD8 | BV786 | RPA-T8 | 563823 | BD Biosciences |
| NKG2A | PE-Vio770 | REA110 | 130-113-567 | Miltenyi |
| NKG2C | PE | 134591 | FAB138P | R&D Systems |
| NKG2C | PE | REA205 | 130-119-776 | Miltenyi |
| KIR2DL1/S1 | FITC | REA284 | 130-118-961 | Miltenyi |
| KIR2DL2/L3/S2 | FITC | CH-L | 559784 | BD Biosciences |
| KIR3DL1/S1 | FITC | REA168 | 130-104-836 | Miltenyi |
| NKG2C | Biotin | REA205 | 130-120-448 | Miltenyi |
| mCD45 | BV570 | 30-F11 | 103135 | BioLegend |
| mCD11b | BV605 | M1/70 | 101237 | Biolegend |
| mF4/80 | BV510 | BM8 | 123135 | Biolegend |
| mCD14 | BV785 | Sa14-2 | 123337 | Biolegend |
| mLy6C | AF700 | HK1.4 | 128023 | BioLegend |
| mCD193 | BV421 | J073E5 | 144517 | Biolegend |

| | | | | |
|---|---|---|---|---|
| mCD206 | BV650 | C068C2 | 141723 | Biolegend |
| mCD11c | BV711 | N418 | 117349 | Biolegend |
| iNOS | PE | W16030C | 696805 | Biolegend |
| mSiglec F | PE Dazzle 594 | S17007L | 155529 | Biolegend |
| mLy6G | PE-Cy7 | 1A8 | 127617 | BioLegend |
| mMHC II | APC-Fire 750 | M5/114.15.2 | 107651 | Biolegend |
| mArginase-1 | APC | A1exF5 | 17-3697-80 | Thermo Scientific |
| LD Green | | | L34970 | Thermo Scientific |
| LD Aqua | | | L34966 | Thermo Scientific |
| DAPI | | | 564907 | BD Biosciences |
| Human FC Blocking Reagent | | | 130-059-901 | Miltenyi |
| Anti-Mouse FC Block | | | 553142 | BD Biosciences |
| CBA Mouse Mig Flex Set | | | 558341 | BD Biosciences |
| CBA Mouse GM-CSF Flex Set | | | 558347 | BD Biosciences |
| CBA Mouse G-CSF Flex Set | | | 560152 | BD Biosciences |
| CBA Mouse MCP-1 Flex Set | | | 558342 | BD Biosciences |
| CBA Mouse IL-6 Flex Set | | | 558301 | BD Biosciences |
| CBA Mouse IL-3 Flex Set | | | 558346 | BD Biosciences |
| CBA Mouse IL-5 Flex Set | | | 558302 | BD Biosciences |
| CBA Mouse IFN-y Flex Set | | | 558296 | BD Biosciences |
| KC/CXCL1 Mouse ELISA Kit | | | EMCXCL1 | Thermo Scientific |
| Mouse SAA-3 ELISA | | | EZMSAA3-12K | Millipore Sigma |
| IL-15/IL-15R Complex Mouse Uncoated ELISA Kit with Plates | | | 88-7215-22 | Thermo Scientific |
| CBA Human GM-CSF Flex Set | | | 558335 | BD Biosciences |
| CBA Human G-CSF Flex Set | | | 558326 | BD Biosciences |
| CBA Human IL-6 Flex Set | | | 558276 | BD Biosciences |
| CBA Human TNF Flex Set | | | 558273 | BD Biosciences |
| CBA Human IL-3 Flex Set | | | 558355 | BD Biosciences |

| | | | | |
|---|---|---|---|---|
| CBA Human IL-2 Flex Set | | | 558270 | BD Biosciences |
| CBA Human IFN-y Flex Set | | | 558269 | BD Biosciences |
| CBA Human Soluble Protein Master Buffer Kit | | | 558264 | BD Biosciences |
| CBA Mouse/Rat Soluble Protein Master Buffer Kit | | | 558266 | BD Biosciences |
| NK Cell Isolation Kit, Human | | | 130-092-657 | Miltenyi |
| Anti-Biotin UltraPure Microbeads | | | 130-105-637 | Miltenyi |
| CD8 microbeads, Human | | | 130-045-201 | Miltenyi |
